# Binding Site Maturation Modulated by Molecular Density Underlies Ndc80 Binding to Kinetochore Receptor CENP-T

**DOI:** 10.1101/2024.02.25.581584

**Authors:** Ekaterina V. Tarasovetc, Gunter B. Sissoko, Aleksandr Maiorov, Anna S. Mukhina, Fazoil I. Ataullakhanov, Iain M. Cheeseman, Ekaterina L. Grishchuk

## Abstract

Macromolecular assembly depends on tightly regulated pairwise binding interactions that are selectively favored at assembly sites while being disfavored in the soluble phase. This selective control can arise due to molecular density-enhanced binding, as recently found for the kinetochore scaffold protein CENP-T. When clustered, CENP-T recruits markedly more Ndc80 complexes than its monomeric counterpart, but the underlying molecular basis remains elusive. Here, we use quantitative *in vitro* assays to reveal two distinct mechanisms driving this behavior. First, Ndc80 binding to CENP-T is a two-step process: initially, Ndc80 molecules rapidly associate and dissociate from disordered N-terminal binding sites on CENP-T. Over time, these sites undergo maturation, resulting in stronger Ndc80 retention. Second, we find that this maturation transition is regulated by a kinetic barrier that is sensitive to the molecular environment. In the soluble phase, binding site maturation is slow, but within CENP-T clusters, this process is markedly accelerated. Notably, the two Ndc80 binding sites in human CENP-T exhibit distinct maturation rates and environmental sensitivities, which correlate with their different amino-acid content and predicted binding conformations. This clustering-induced maturation is evident in dividing human cells, suggesting a distinct regulatory entry point for controlling kinetochore assembly. We propose that the tunable acceleration of binding site maturation by molecular crowding may represent a general mechanism for promoting the formation of macromolecular structures.

**Significance Statement:** A distinctive mechanism of protein-protein interaction underpins the assembly of kinetochores, which is critical for human cell division. During mitosis, the Ndc80 complex must bind tightly to the unstructured N-terminus of its receptor, CENP-T, which is densely clustered at kinetochores. Using single-molecule *in vitro* assays, we show that Ndc80 binding is mediated by an initially unstable yet tunable interface. The high molecular density of CENP-T at the kinetochores accelerates the maturation of this binding interface, favoring the formation of stable complexes within the kinetochore structure, rather than in the soluble phase. This environment-driven modulation of binding site maturation may represent a key regulatory mechanism for ensuring strong and specific interactions during the assembly of macromolecular complexes such as kinetochores.

## Introduction

During the biogenesis of higher-order cellular assemblies, different proteins must come together from the cytosol to build robust structures on biologically-relevant timescales. However, how the binding of individual components is directed specifically to the assembling structures, while preventing the same components from forming potentially toxic and wasteful complexes in the cytosol, is not well understood. In dividing cells, the assembly of mitotic kinetochores involves a hierarchical array of pairwise protein-protein interactions. Among these assembly steps is the recruitment of the core kinetochore Ndc80 complex from its 50-150 nM cytosolic pool to the kinetochore-localized CENP-T receptor protein, which is also present in a cytosol at 5-25 nM (1-4). Several known mechanisms regulate Ndc80 recruitment by CENP-T, including mitotic-specific CENP-T phosphorylation (5-12), nuclear-cytoplasmic compartmentalization during interphase (5) and activation of the full length Ndc80 complex (13). However, these regulatory mechanisms fail to explain our recent findings in mitotic human cells that formation of Ndc80/CENP-T complexes depends on their specific molecular environment (14). Indeed, the disordered N-terminus of CENP-T, which contains two Ndc80 binding sites, recruits Ndc80 poorly in mitotic cytoplasm, suggesting weak Ndc80 binding affinity (6, 13-15). By contrast, the artificially-generated CENP-T oligomers expressed at the same protein level as the monomeric CENP-T recruit Ndc80 efficiently, implying that within the kinetochore-mimicking clusters CENP-T and Ndc80 have stronger binding affinity (14). Because CENP-T molecules are present in multiple copies at mitotic kinetochores (16), the molecular density-dependent recruitment of Ndc80 molecules is of critical physiologically importance (14), necessitating a detailed mechanistic investigation of this apparent affinity paradox. Here, we determined the kinetics of Ndc80/CENP-T interactions *in vitro* to reveal an elegant regulatory mechanism involving a molecular density-dependent kinetic barrier for maturation of the Ndc80/CENP-T binding interfaces.

## Results

### The interaction between Ndc80 and CENP-T does not follow a straightforward single-step reaction mechanism

We first analyzed the behavior of a monomeric CENP-T by immobilizing its N-terminal fragments at low density on a coverslip in a flow chamber (Fig. 1A-C, *SI Appendix*, Fig. S1). Using total internal reflection fluorescence (TIRF) microscopy, we monitored binding of soluble GFP-tagged “Bonsai” Ndc80 complex with an internal truncation (17), thereafter called Ndc80 complex). At 200 nM Ndc80 complex, binding of Ndc80 to monomers of CENP-T^6D^ activated with phosphomimetic substitutions quickly plateaued at roughly two Ndc80 molecules (Fig. 1D). Thus, at Ndc80 concentrations comparable to intracellular levels (1, 2), both binding sites become saturated and achieve steady-state within ∼1 min, indicating fast association. Binding of the GFP and its nanobody plateaued at 0.9, which is close to the expected 1:1 ratio when all binding sites are accessible (Fig. 1D). This assay did not detect binding of the Ndc80 construct lacking its CENP-T binding domain Spc24/25 (Fig. 1D), confirming the specificity of these interactions.

**Figure 1.**
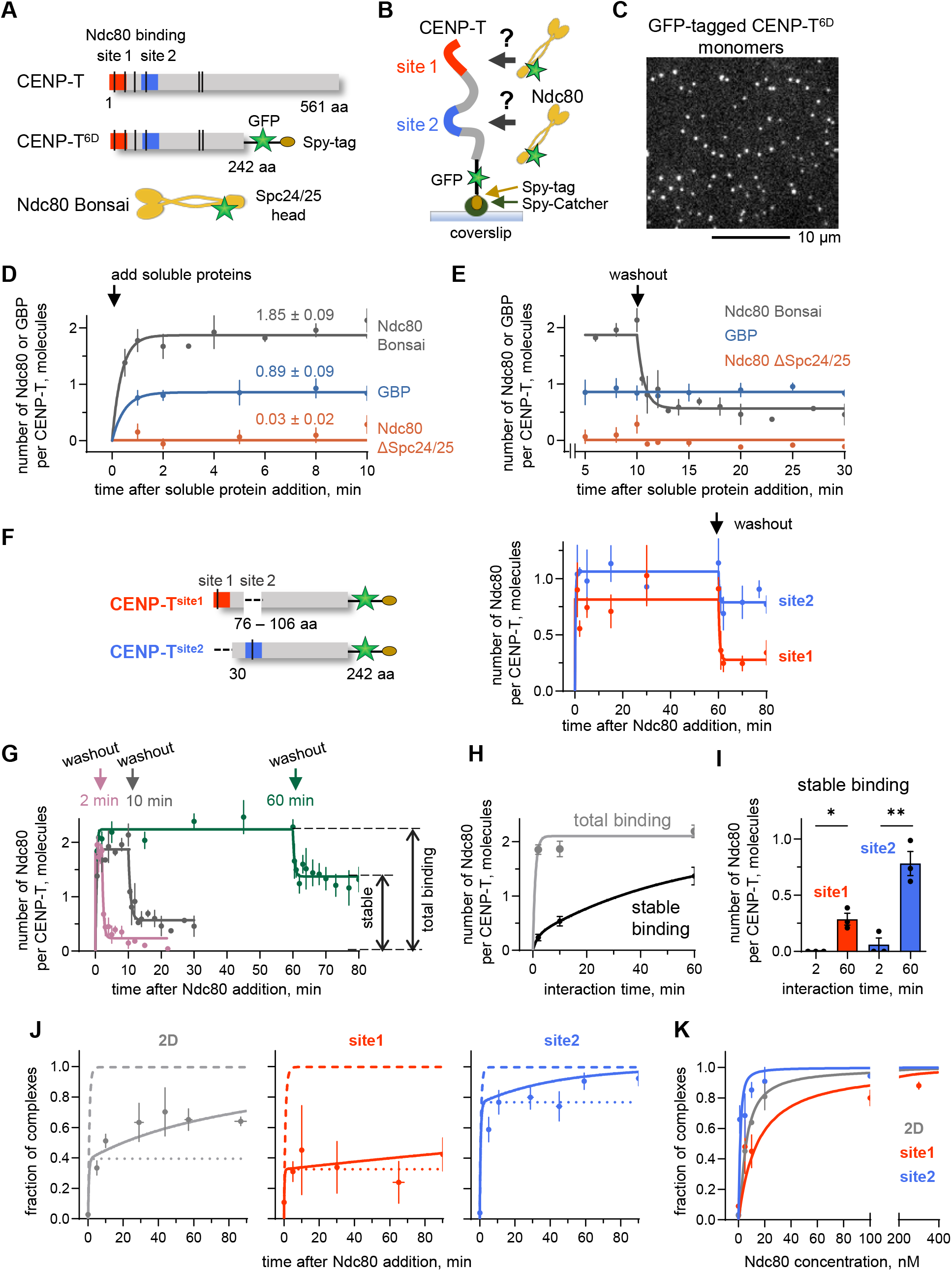
Ndc80 association with monomeric CENP-T and Ndc80/CENP-T complex stability. **(A)** Diagram of human CENP-T protein and CENP-T^6D^ construct with GFP tag for visualization, and Spy-tag for immobilization. **(B)** Schematics of the experiment with the coverslip-immobilized CENP-T^6D^ monomers. **(C)** Example imaging field. **(D)** Binding of indicated fluorescent proteins (200 nM) to monomeric CENP-T^6D^. Numbers are the steady-state levels with SEM, lines are exponential fittings. Here and in other panels, each point represents mean ± SEM for N = 3-8 independent experiments each imaging > 20 molecules, see Source Data file. **(E)** Dissociation of indicated proteins from CENP-T^6D^ monomers after removal of soluble proteins. **(F)** Diagram of the CENP-T^site1^ (site1) and CENP-T^site2^ (site2) constructs, and binding and dissociation of 200 nM Ndc80 for coverslip-immobilized CENP-T monomers. **(G)** Binding of 200 nM Ndc80 to CENP-T^6D^ monomers and dissociation kinetics for different incubation times. **(H)** Time-dependent changes in the number of total (in the presence of soluble Ndc80) and stable (after Ndc80 washout) Ndc80 molecules per CENP-T^6D^ monomer. **(I)** Number of the stably bound Ndc80 molecules. Bars show mean ± SEM; each point corresponds to an independent experiment with > 20 molecules; unpaired t-test with Welch’s correction: * = p < 0.05, ** = p < 0.01. **(J)** Kinetics of complex formation for 10 nM Ndc80 and 1 nM CENP-T proteins studied with FCS. Points are experimental data (mean ± SEM, N = 2-5). For CENP-T^2D^, fraction of complexes with two Ndc80 molecules is plotted. Lines are predictions of different models: solid line – model with site maturation, broken lines – conventional binding with different dissociation constants, see *SI Appendix*, Note 2. **(K)** Fraction of Ndc80/CENP-T complexes in FCS experiments with 1 nM CENP-T: experimental points and curves predicted by the model with site maturation.

To probe the stability of Ndc80/CENP-T complexes, we abruptly removed soluble Ndc80 by washout (Fig. 1E). Surprisingly, the Ndc80 dissociation kinetics were bi-phasic, with most Ndc80 detaching immediately, but ∼30% remaining as a stable population. In contrast, complexes between GFP and its nanobody were highly stable and did not undergo dissociation. Because the CENP-T has two binding sites for Ndc80 (6, 15), we tested whether they had different dissociation rates. We generated deletion constructs in which only one of the two sites was preserved (Fig. 1F). Each site quickly bound ∼1 Ndc80 molecule and, although they retained different fractions of stable molecules, the Ndc80 dissociation upon washout was bi-phasic for each site (Fig. 1F). The lack of the single-step dissociation kinetics for these protein complexes strongly implies that the interaction between Ndc80 and each of its two binding sites on CENP-T is not a straightforward, single-step binding reaction, and it involves additional layers of complexity or multiple stages.

### Initial Ndc80 binding by CENP-T is unstable, but Ndc80 retention increases over time

To gain insight into the underlying reaction mechanism, we varied the incubation time of Ndc80 and CENP-T^6D^ prior to a washout. In a single-step reaction, removing Ndc80 at any time after the CENP-T sites have saturated and the binding has reached steady-state should result in an identical, time-independent outcome. Strikingly, the fraction of stably-bound Ndc80 increased progressively with the incubation time (Fig. 1G). The time-dependent increase in complex stability could not be attributed to the increase in total protein binding during longer incubations, because the total number of bound Ndc80 molecules remained close to 2 during the entire experiment (Fig. 1H). The increased Ndc80 retention also could not be explained by changes in CENP-T alone, because the stability of Ndc80/CENP-T complex remained unchanged after the CENP-T was incubated with a buffer alone for an extended time prior to Ndc80 addition (*SI Appendix*, Fig. S2A, Note 1.1). Continuous imaging of Ndc80/CENP-T complexes in the presence of soluble Ndc80 also revealed time-dependent changes in complex stability (*SI Appendix*, Note1.2, Fig. S2B), so this effect does not depend on Ndc80 removal by a washout.

Moreover, the time-dependent stabilization of the Ndc80/CENP-T complexes was observed for each of the CENP-T binding sites (Fig. 1I; *SI Appendix*, Fig. S2C). To test whether the two-step character of Ndc80 and CENP-T interaction is caused by their proximity to the surface of the coverslip, we studied their binding in solution using Fluorescence Correlation Spectroscopy (FCS) (*SI Appendix*, Note 1.3, Fig. S3). Binding between Ndc80 and CENP-T proteins containing both or just one of the sites showed bi-phasic kinetics, strongly supporting the two-step interaction mechanism (Fig. 1J,K). Thus, although CENP-T sites bind to Ndc80 rapidly, an extended interaction time is needed to develop the high-affinity bonding, a property that we refer to as “maturation”. Curiously, under the tested conditions using physiological Ndc80 concentrations with CENP-T monomers, the stable retention of both Ndc80 molecules was not achieved even after 60 min, which exceeds the time of chromosome segregation in human cells suggesting the need for additional stabilization mechanisms.

### Clustering of CENP-T does not affect the initial fast binding of Ndc80 and the biphasic character of dissociation

In mitotic cells, interactions between soluble CENP-T and Ndc80 components are weak, whereas CENP-T scaffolds at the kinetochore form high-affinity, load-bearing bonds with Ndc80 proteins (14, 18, 19). Prior work proposed that clustering of CENP-T scaffolds at kinetochores promotes Ndc80 recruitment in mitotic cells, based on findings that Ndc80 binding is enhanced when CENP-T is multimerized in the cytoplasm and *in vitro* (14). However, the mechanistic basis for this enhancement remains undetermined. The improved Ndc80 recruitment in the high-density environment may result from an increased association rate with CENP-T or a decreased dissociation rate of the Ndc80/CENP-T complex. Since Ndc80 binding to CENP-T does not occur through a single-step reaction, the recruitment may also depend on the transition rate from a low-affinity to a high-affinity state. To distinguish between these non-mutually exclusive possibilities, we used our quantitative TIRF assay with CENP-T clusters.

CENP-T^6D^ proteins were conjugated to a 60-subunit dodecahedral “mi3” particle (20, 21) to achieve clustering of 42 ± 2 molecules (Fig. 2A; *SI Appendix*, Fig. S4, S5), which approximates the ∼70 copies of CENP-Ts at the human kinetochore (16). Similar to monomeric CENP-T, our time-resolved binding assay showed that up to 2 Ndc80 molecules rapidly and specifically associated with each clustered CENP-T molecule (Fig. 2B,C). Incubation with Ndc80 lacking the Spc24/25 domains showed no binding, and no or minimal binding was observed between Ndc80 Bonsai and clusters containing a fragment of CENP-T lacking the Ndc80 binding sites (106-242 aa) or Ndc80 Bonsai alone (*SI Appendix*, Fig. S6 A-C), confirming that these interactions rely on the conventional Ndc80/CENP-T binding interfaces. After saturation of both sites on the clustered CENP-T^6D^ proteins, soluble Ndc80 was removed. The fraction of stably bound molecules increased significantly in clusters of CENP-T^6D^, as well as CENP-T proteins with only one of the binding sites relative to their respective monomers (*SI Appendix*, Fig. S6D). Importantly, the overall shape of the dissociation curves was highly similar for clustered and monomeric CENP-T: the initial dissociation was fast, followed by a plateau with the stably-bound complexes, albeit at different levels (Fig. 2C). Thus, the kinetics of initial Ndc80 binding and the dissociation rates for the two populations of Ndc80/CENP-T complexes are maintained in clustered CENP-T and these kinetic features are not responsible for the increased fraction of stably bound Ndc80 molecules.

**Figure 2.**
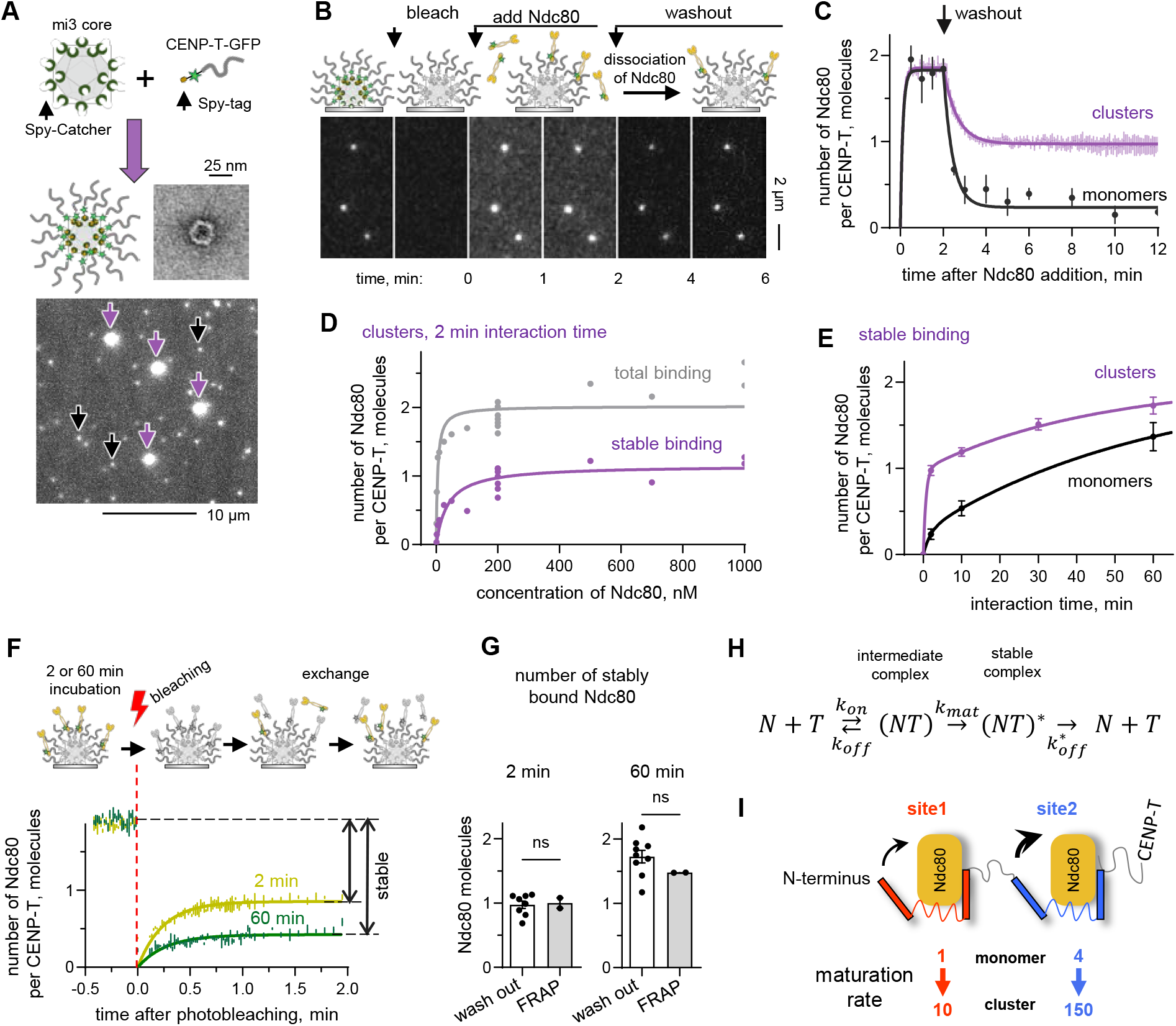
Ndc80 association with clustered CENP-T and Ndc80/CENP-T complex stability. CENP-T multimerization strategy, an electron microscopy image of an assembled cluster, and representative GFP-fluorescence image of the coverslip-immobilized GFP-tagged CENP-T^6D^ monomers (black arrows) and their mi3-based clusters (purple). **(B)** Schematics of the experiment and representative images of GFP-tagged CENP-T^6D^ clusters; unless indicated otherwise, Ndc80-GFP was 200 nM. Bleaching of GFP-CENP-T clusters prior to Ndc80-GFP addition was used to discriminate their corresponding signals and enable ratiometric measurements. **(C)** Binding kinetics and dissociation for CENP-T^6D^ molecules; results for monomers are same as in Fig. 1G. **(D)** Number of total and stable Ndc80 molecules per one CENP-T^6D^ in clusters; each point is an independent experiment with > 12 clusters; lines are hyperbolic fits. **(E)** Kinetics of stable complex formation between Ndc80 and CENP-T^6D^ molecules in clustered vs. monomeric (same as in Fig. 1H) form. **(F)** Schematic of the FRAP experiment to assess Ndc80 exchange at steady-states. CENP-T^6D^ clusters were preincubated with Ndc80 for 2 or 60 minutes yielding two bound Ndc80 molecules but different fraction of non-exchanging complexes. **(G)** Stably bound molecules with each point representing one independent experiments; bars show mean ± SEM. The lack of significance (ns) was determined using unpaired t-test. **(H)** Two-step reaction scheme, see *SI Appendix*, Note 2 for details. **(I)**. The diagram summarizing quantitative findings for maturation of sites in monomeric vs. clustered CENP-T. Ndc80 is shown binding to two interfaces of each CENP-T site, symbolizing intermediate complex. Maturation transition is depicted with the curved black arrows that bring the third interface in contact with Ndc80, thereby increasing the binding strength. The numbers are normalized maturation rates relative to the slowest transition (site 1 in CENP-T^2D^ monomers), which was 2 × 10^−4^ s^-1^.

### Clustering of CENP-T accelerates Ndc80 retention

The biphasic Ndc80 dissociation from the CENP-T^6D^ clusters implies that interactions between Ndc80 and clustered CENP-T proceed by a similar mechanism as with CENP-T monomers. To test this, we increased Ndc80 concentration up to 1 μM. However, despite this saturating level of Ndc80, the outcome did not change when incubation time was kept constant (2 min, Fig. 2D). Two Ndc80 molecules were still bound in the presence of soluble Ndc80, and the fraction of stable complexes did not increase, consistent with the fast initial Ndc80 binding being a straightforward one-step reaction. However, increasing the incubation time between CENP-T clusters and 200 nM Ndc80 up to 60 min led to a significantly increased fraction of the strongly bound Ndc80 molecules, similar to CENP-T monomers (Fig. 2E).

To test whether the time-dependent stabilization of the Ndc80/CENP-T complexes was affected by our specific assay employing soluble protein removal, we measured the recovery of Ndc80-GFP binding to non-fluorescent CENP-T clusters after photobleaching (Fig. 2F). This FRAP assay carried out in the presence of soluble Ndc80-GFP showed a highly similar increase in Ndc80 retention by the CENP-T clusters (Fig. 2G). Thus, maturation of the Ndc80/CENP-T complexes is a bona-fide feature of both monomeric and clustered molecules. The substantially larger fraction of the stably-bound Ndc80 per CENP-T in clusters vs. monomers corresponding to the high-affinity state is achieved at a faster rate in a denser molecular environment, where multiple and diverse molecular contacts may affect conformational dynamics of interacting proteins.

To estimate the underlying rate constants, we constructed a mathematical model based on the proposed reaction scheme (Fig. 2H; *SI Appendix*, Note 2, Fig. S7). A wide range of our experiments using different CENP-T proteins could be described using the same or slightly adjusted binding/unbinding rate constants (*SI Appendix*, Table S1). This includes the binding/unbinding curves for different CENP-T clusters (CENP-T^2D^ and proteins containing single sites) at varying incubation times and Ndc80 concentrations (*SI Appendix*, Fig. S8), as well as our experimental results for coverslip-immobilized CENP-T monomers (*SI Appendix*, Fig. S9) and soluble molecules (Fig. 1J,K). This consistency across diverse experimental approaches provides confidence in the conclusions drawn from our data. In particular, these quantitative analyses revealed that the kinetic rates for the intermediate complexes with Ndc80 at the two binding sites in CENP-T are similar, with site 2 having a 3-fold faster association rate and a ∼6-fold slower dissociation rate. The resulting ∼10-fold difference in the affinities of these intermediate complexes, however, does not account for the observed bi-phasic kinetics, which arises from the transition into a stable complex state. Although experiments with different CENP-T clusters could be well described using the same binding/unbinding rate constants as for their respective monomers, for all proteins and experiments, the maturation rates that applied to the monomers had to be significantly increased for CENP-T clusters. In CENP-T^2D^ clusters, the maturation rate increased nearly 40-fold for site 2 and 10-fold for site 1 (Fig. 2I). Thus, both binding sites are highly sensitive to their molecular environment, with site 2 being particularly responsive.

### The rate of Ndc80/CENP-T complex stabilization correlates with the sequence and conformation of the CENP-T binding site

The enhanced retention of Ndc80 after forming an intermediate complex with CENP-T suggests a transformation of their binding interface that strengthens affinity. To investigate the molecular determinants of this transition, we analyzed the properties of two binding sites. The Ndc80-binding interface on CENP-T involves a ∼30-amino-acid linear stretch, predicted to contain a central α-helix flanked by unstructured regions (Fig. 3A; *SI Appendix*, Fig. S10, S11). In the published structure of chicken proteins, a CENP-T peptide with this tri-partite organization wraps tightly around the Spc24/25 globular domain (22). In human CENP-T, AlphaFold3 predicts similar positions for central α-helices within the Spc24/25 groove, although these helices differ by four amino acids, altering their charge and hydrophobicity (Fig. 3B; *SI Appendix*, Note 3) (23). The predictions for the flanking regions for site 1 have low confidence (*SI Appendix*, Fig. S11, Movie 1), indicating that their conformations may vary over time or depending on the molecular environment. For site 2, while conformation of the N-terminal flank is also variable, the C-terminal flank reproducibly forms a more extensive interface with Spc24/25 compared to site 1 (Fig. 3B; *SI Appendix* Fig. S12, Movie 2). In our *in vitro* assays, site 1 exhibited slower maturation than site 2 (Fig. 2I), suggesting that conformational changes in the unstructured flanking regions may influence maturation. However, differences in the molecular makeup of the central α-helices, or the distinct positional locations of the two sites within CENP-T, with site 1 being at the N-terminus, may also contribute to the observed differences in maturation rates.

**Figure 3.**
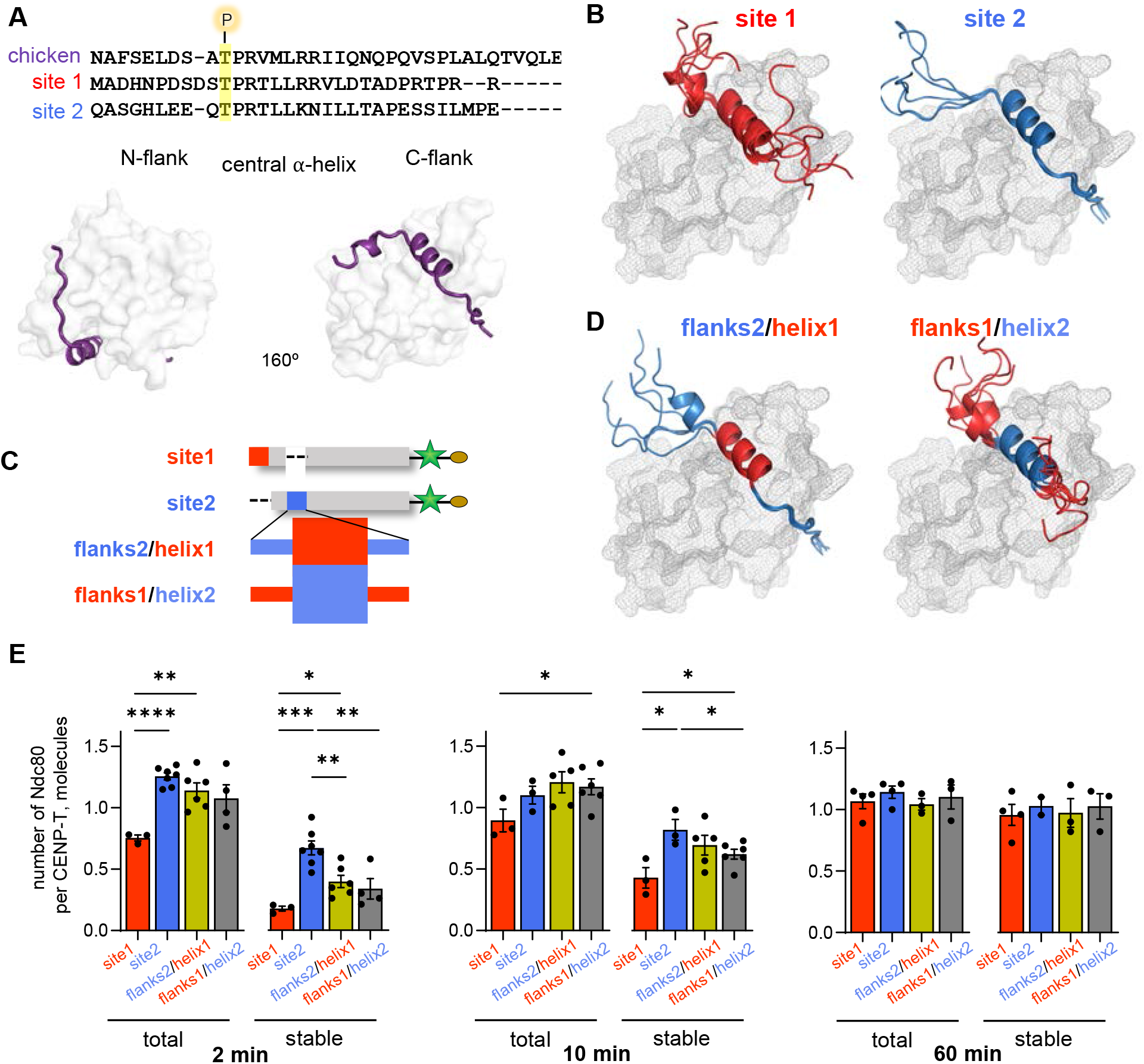
Molecular analysis of Ndc80/CENP-T binding interface. **(A)** Sequence alignment of chicken and two human Ndc80-binding sites with conserved phosphorylated threonine. Structure of chicken complex (PDB: 3VZA) illustrating the tri-partite binding interface with the central α-helix inserted into a Spc24/25 groove, and two flanking regions wrapped around the Spc24/25 head. (**B**). AlphaFold3-based predictions (N = 5) for human CENP-T sites in complex with Spc24/25 globule of the Ndc80 complex, shown as grey surfaces. **(C)** Diagram of CENP-T with mutant site 2 in the CENP-T protein lacking site 1. **(D**) Same as in panel (B) but for composite site 2. **(E)** Total and stable binding of Ndc80 (200 nM) to indicated CENP-T clusters. Each point represents an independent experiment with > 13 clusters; bars are mean ± SEM; unpaired t-test: * = p < 0.05, ** = p < 0.01, *** = p<0.001, **** = p<0.0001; data sets with p > 0.05 are not labeled.

To investigate the relative contributions of these molecular features, we engineered mutant CENP-T proteins in which site 1 was removed and its central helix or flanking regions were inserted into the corresponding positions of site 2 (Fig. 3C). Based on AlphaFold3 predictions, replacing the flanks resulted in variable conformations at both flanks, mimicking site 1, while substituting helix 2 with helix 1 preserved the more ordered positioning of the C-terminal flank of site 2 (Fig. 3D, Movies 3,4). We directly tested the effects of these perturbations by monitoring Ndc80 binding and dissociation in clusters formed by these mutant proteins. Both mutants retained normal intermediate complex formation on par with the original site 2, as indicated by the rapid binding of one Ndc80 molecule per CENP-T (Fig. 3E). Additionally, they preserved the ability to mature, with stable retention of one Ndc80 molecule after 60 minutes. However, the maturation rate of site 2 was significantly reduced when its various regions were replaced with the corresponding sequences from site 1, suggesting that the location of site 2 alone cannot account for its faster maturation. Both mutants exhibited a similar reduction in maturation rate, implying that no single distinguishing feature, such as the degree of disorder in the C-terminal flank, can fully explain the differences between the two sites. Thus, the maturation rate of the tri-partite binding interface is influenced by both the composition of the central helix and the flanking regions, likely reflecting their complex dynamic behavior.

### Ndc80 binding sites on CENP-T exhibit differential clustering-dependent regulation in mitotic cells

Our *in vitro* assays revealed that the maturation of site 2 was accelerated more strongly in a dense molecular environment, whereas maturation of site 1 was less sensitive to CENP-T clustering (Fig. 2I). Next, we tested if the different maturation rate of the two sites is preserved in dividing cells and leads to phenotypic differences. Since it is challenging to distinguish the recruitment of Ndc80 via different sites and pathways at intact kinetochores, we took advantage of our sensitized outer kinetochore assembly assay (14). We fused a 242 amino acid N-terminal CENP-T to a single-chain monoclonal antibody (scFv), and co-expressed it in HeLa cells with a tdTomato-tagged scaffold that contained multiple repeats of the antibody’s cognate epitope, GCN4pep (24) (Fig. 4A). By varying the number of GCN4pep repeats, we formed multimers consisting of 1-12 CENP-T molecules with different sequences. Expression of unclustered CENP-T^WT^ did not result in cellular consequences, with a similar fraction of mitotic cells to uninduced control cells (Fig. 4B; *SI Appendix*, Fig. S13A). However, increasing CENP-T multivalency by increasing the number of GCN4pep repeats on the scaffold in cells expressing the same level of CENP-T^WT^ protein led to a pronounced mitotic arrest, indicating assembly of cell-cycle disrupting kinetochore-like particles consistently with prior report (14). To investigate clustering-dependent behavior of two sites, we generated CENP-T constructs with only one active Ndc80 site by inactivating the other site with an alanine substitution. CENP-T multimers with deactivated site 1 (CENP-T^T11A^) mirrored the clustering-dependency of CENP-T^WT^ (Fig. 4B). Conversely, the CENP-T mutant with deactivated site 2 (CENP-T^T85A^) exhibited reduced potency, aligning with our expectations from *in vitro* experiments.

**Figure 4.**
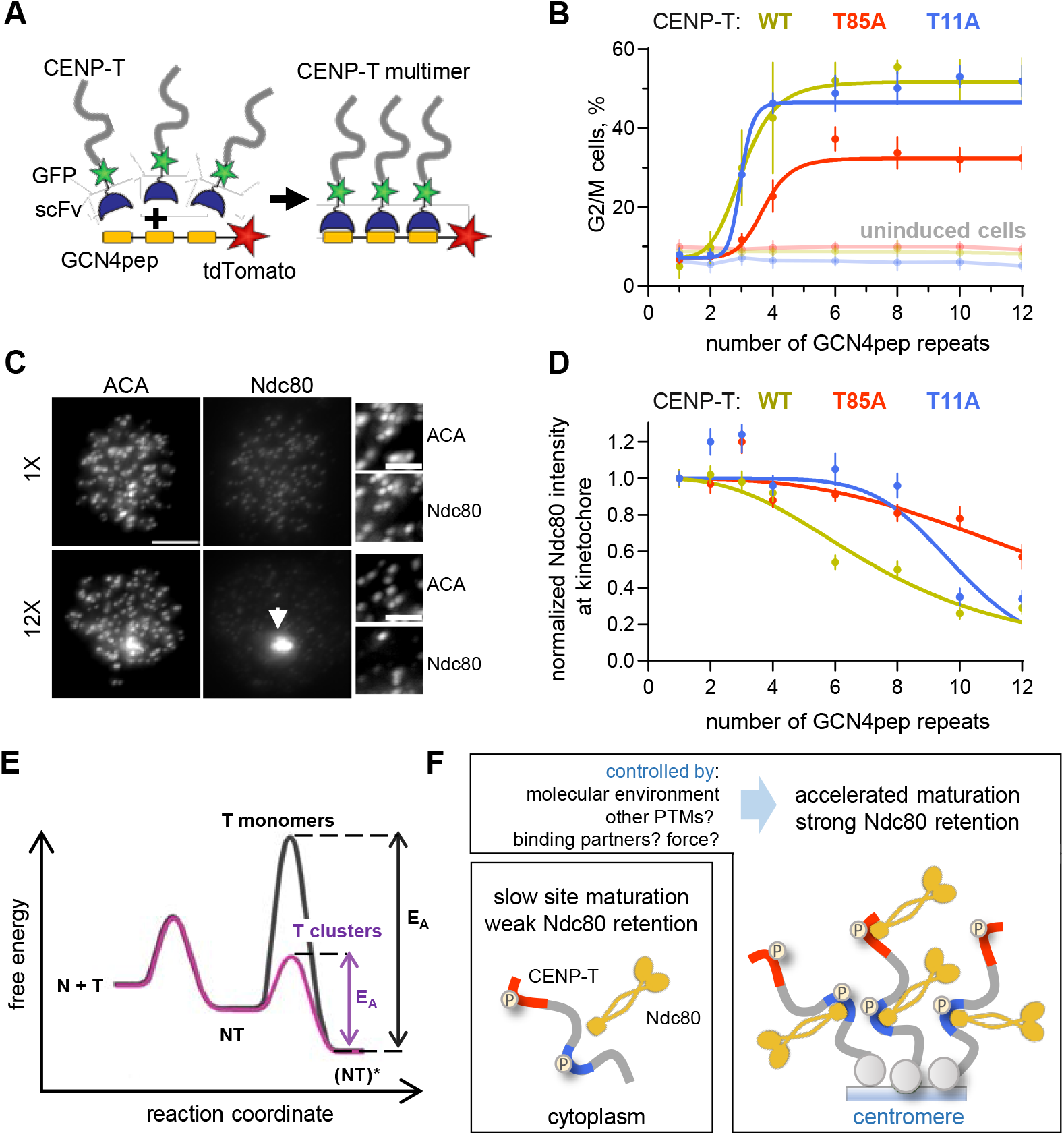
Distinct clustering-dependent regulation of Ndc80 binding sites in mitotic cells. **(A)** Diagram illustrating the sensitized outer kinetochore assembly assay utilizing Sun-Tag tunable oligomerization. **(B)** Percent of G2/M cells for cells expressing CENP-T constructs oligomerized using Sun-Tag scaffolds with different number of GCN4pep. Data for uninduced cells are shown in faded colors. Each point represents the mean ± SEM, N = 2-4, lines are fits with a sigmoidal function. Data in panel (B) and (D) for CENP-T^WT^ construct are from (14), and are provided here for a direct comparison. **(C)** Representative images of mitotic cells showing localization of kinetochores and various CENP-T^WT^ oligomers (white arrows). The centromeres (ACA) and Ndc80 are visualized with immunofluorescence. Insets are adjusted differently from the full-size images for improved visibility. Scale bar: 5 μm. **(D)** Same experiments as in panel (B) but showing Ndc80 kinetochore level; 23-46 total cells for each condition pooled from N = 3 for CENP-T^WT^ and N = 2 for CENP-T^T11A^ and CENP-T^T85A^. **(E)** Free energy difference (not to scale) between the unbound and complexed Ndc80 (N) and CENP-T (T) proteins plotted against the reaction coordinate. NT – intermediate complex. Activation energy barrier (EA) for a strong-affinity state (NT)* is sensitive to the molecular density of CENP-T. **(F)** Simplified schematics of the established and proposed mechanisms to control direct Ndc80 recruitment by CENP-T scaffold. For simplicity, shortened molecular variants are depicted and other kinetochore proteins are omitted.

To confirm that the mitosis-specific toxicity of various CENP-T multimers correlated with their ability to recruit Ndc80 and thereby compete with endogenous kinetochores (14), we quantified kinetochore-localized Ndc80 levels. CENP-T multimers with wild type CENP-T sequences resulted in a substantial reduction in kinetochore-localized Ndc80 across a range of multimer copy numbers, effectively competing with the endogenous CENP-T receptors. As the size of cytoplasmic CENP-T^WT^ multimers increased, kinetochores gradually lost Ndc80 down to 20%, concomitant with the formation of Ndc80-containing particle aggregates (Fig. 4C,D; *SI Appendix*, S13B). CENP-T^T11A^, containing only active site 2, exhibited an Ndc80-stripping effect similarly to CENP-T^WT^, although at a slightly larger clustering size. In contrast, CENP-T^T85A^, containing active site 1, showed less activity, resulting only in a 60% reduction of kinetochore-bound Ndc80 in the presence of CENP-T 12-mers. These results demonstrate that site 2 is largely responsible for the behavior of wild type CENP-T, displaying a stronger activation by CENP-T oligomer size than site 1. Thus, the different maturation rates observed *in vitro* for the two CENP-T binding sites correlate with their differential sensitivity to molecular density in mitotic cells.

## Discussion

Understanding biophysical principles that enable the assembly of higher-order structures, like those represented by the mitotic kinetochores, is fundamentally important and has strong potential for applications in medicine and nanoengineering (25, 26). Here, we investigated the kinetic mechanisms of a critical assembly reaction at human kinetochores, in which molecules of the core component Ndc80 are linked to dozens of CENP-T receptors within the dense molecular meshwork (16). Ndc80/CENP-T complexes sustain forces during chromosome segregation, implying that they are linked via the high-affinity interface that is not reversible at physiological timescale (18, 19). Thus, specific molecular mechanisms must ensure that the Ndc80 and CENP-T avoid forming such strong interfaces in a cytosol, while bonding strongly at the kinetochore.

Our study reveals that this recruitment specificity can be mediated by two interconnected mechanisms. The first mechanism involves the newly discovered ability of a weak Ndc80/CENP-T interface to mature into a high affinity state, whereas the second mechanism features acceleration of this transition by the molecular environment for these interactions.

The interface maturation was revealed by our finding that binding of Ndc80 molecules to their site on CENP-T proceeds via a two-step reaction (Fig. 4E). An intermediate Ndc80/CENP-T complex with low affinity forms rapidly. However, a different binding mode — exceeding two orders of magnitude in affinity — is also possible. This transition likely involves conformational changes in the unstructured flanking regions of the CENP-T binding site, though the central α-helix composition also plays a crucial role. Spontaneous formation of the full, topologically complex interface in which the flanking regions wrap around the Spc24/25 domain may be kinetically challenging. Indeed, the binding kinetics of proteins with unstructured regions often involve multiple steps (27, 28). Resolving the precise conformational dynamics at the Ndc80 and CENP-T binding interface is a complex task that will require extensive future research.

To ensure the selective formation of the Ndc80/CENP-T complex specifically at the higher-order structures, the kinetic parameters of their molecular interactions must be sensitive to the exact molecular environment. Prior work has demonstrated that dense molecular environment formed by proteins with low-complexity regions, such as within liquid condensates, can modulate molecular interactions and kinetic steps (29, 30). By comparing Ndc80 interaction with CENP-T clusters vs. monomers, we found little difference in the rate of the intermediate complex formation and its stability, as well as in the stability of the high-affinity state (*SI Appendix*, Table S1). However, the rate of the stable complex formation was remarkably sensitive to CENP-T clustering. Dense molecular environment may accelerate conformational dynamics of the flanking unstructured regions of the CENP-T site either by steric or weak multivalent interactions with neighboring molecules. Indeed, densely crowded environment within molecular condensates can enhance protein activity by promoting specific interactions and facilitating enzymatic reactions (29, 30), suggesting that activation of the CENP-T site within the clusters may rely on similar mechanisms. We propose that this density-dependent acceleration of the otherwise slow two-step binding reaction is responsible for the previously reported preferential Ndc80 recruitment to CENP-T oligomers in mitotic cells (14).

A gap remains in understanding the details and extent of CENP-T site maturation at mitotic kinetochores. Our *in vitro* findings suggest the existence of kinetochore populations of Ndc80 with different stability. Although this prediction is consistent with the presence of a prominent stable Ndc80 fraction at kinetochores in cells (31-35) and its increase during metaphase (33, 35), future work should examine the dynamics of Ndc80 populations in human cells under less intrusive conditions, such as with normal protein levels and with fully functional fluorescent Ndc80 proteins. The physiological significance of the differing maturation rates of two Ndc80 sites on human CENP-T is also presently unknown. This result is particularly intriguing given that CENP-T in some vertebrate organisms, including the well-studied chicken protein, has only one site for Ndc80 binding that appears to be more similar to human site 2 (22). The enhanced sensitivity to CENP-T clustering of human site 2 relative to site 1 has been observed *in vitro* and in intact mitotic cells. We speculate that the tunable interfaces of these sites provide an entry point for additional regulation of kinetochore-microtubule interactions. Previous studies of microtubule attachment and error correction at kinetochores emphasized the controlled binding affinity between Ndc80 complex and microtubules (36). Our findings suggest that kinetochore-microtubule attachments could also be controlled via the tunable affinity of the Ndc80 to CENP-T. Future work should also investigate whether maturation of Ndc80 bond is found for another Ndc80 receptor CENP-C and other kinetochore proteins, or this phenomenon is exclusive to CENP-T.

## Materials and Methods

Detailed description of the materials and methods used in the study can be found in *SI Appendix*, Materials and Methods: Materials and Methods, Cloning, Cell line generation, Cell Culture, Immunofluorescence, DNA content analysis, Protein expression and purification, Assembly of CENP-T clusters in solution, TIRF microscopy assay to study the interactions between monomeric CENP-T and soluble Ndc80, TIRF microscopy assay to study the interactions between clustered CENP-T and soluble Ndc80, FCS assay, Negative Staining Electron Microscopy, Structural analysis of Ndc80/CENP-T complexes using AlphaFold software, and Mathematical modeling.

## Acknowledgments

We are grateful to Drs. Yale Goldman and Him Shweta for guidance and insightful suggestions using FCS. We thank Drs. Tatyana Svitkina and Anil Chougule for guidance and assistance with electron microscopy. We thank Dr. Haitao Li and Tatyana Kogan for assistance with cloning and purifications of CENP-T proteins, Vladimir Demidov for assistance with data analysis, Veronika Balakshina and Polina Soloveva for technical assistance. Dr. Jennifer Deluca and Jeanne Mick (Colorado State University) kindly provided the plasmid with untagged Ndc80 Bonsai and advised on the purification of Ndc80 constructs. We are grateful to Drs. Mark Howarth and Rolle Rahikainen for advice on purifying mi3-based clusters. We also thank Drs. Yale Goldman, Michael Ostap, Ben Black, Michael Lampson, Mathew Good, and their lab members for thoughtful and insightful discussions.

