## Supplementary Information for "Binding Site Maturation Modulated by Molecular Density Underlies Ndc80 Binding to Kinetochore Receptor CENP-T"

**This PDF file includes:**

Materials and Methods

Supplementary text

Note1. Supporting experimental results

Note 2. Mathematical model of Ndc80 interaction with CENP-T

Note 3. Structural analysis of human CENP-T sites bound to Spc24/25 head

Figs. S1 to S13 with Legends

Table S1 with Legend

Legends to Movies

SI References

**Other supporting materials for this manuscript include the following:**

Movies 1-4

Source Data File

### Materials and Methods

#### Cloning

All CENP-T constructs were generated using a pET28-eGFP-Spy-Tag vector. They contained various N-terminal CENP-T fragments followed by a 3xSGGGG linker, eGFP, second 3xSGGGG linker, myc-tag, Spy-Tag, and a 6xHis tag. The CENP-T$^{6D}$ construct was generated by inserting a synthesized cDNA (Genewiz) encoding 1-242 aa of CENP-T containing six phosphomimetic substitutions (T11D, T27D, S47D, T85D, T195D, S201D). CENP-T$^{2D}$ construct was generated by inserting the DNA fragment encoding CENP-T (1-242 aa) from pKG174 (1) and phosphomimetic substitutions T85D and T11D introduced using QuikChange II Site-Directed Mutagenesis Kit (Agilent, 200523). CENP-T$^{2D,short}$ construct was generated by inserting a synthesized cDNA fragment of CENP-T (1-106 aa) containing T11D and T85D. CENP-T$^{site1}$ and CENP-T$^{site2}$ constructs were generated by deleting sequences encoding 76-106 aa and 2-30 aa from CENP-T$^{2D}$ expression vector, correspondingly, using Q5® Site-Directed Mutagenesis Kit (NEB, E0554S). CENP-T$^{\Delta N}$ construct was generated analogously by deleting the sequence encoding 1-106 aa of CENP-T$^{2D}$. Construct CENP-T$^{flanks2/helix1}$ was generated by introducing mutations (K91R, N92R, I93V, and L95D) into the CENP-T$^{site2}$ construct, thereby converting α-helix of site 2 into the α-helix of site 1. Construct CENP-T$^{flanks1/helix2}$ was generated by replacing a fragment of cDNA encoding 76-106 aa of CENP-T$^{site2}$ with a synthetic cDNA fragment (from IDT) encoding amino acids 1-10 of site 1 (the N-terminal flanking region), amino acids 85-95 of site 2 (the central α-helix), and amino acids 21-30 of site 1 (the C-terminal flanking region. To optimize expression and protein purification, the GST-tag with TEV protease cleavage site was inserted at the 5' of CENP-T$^{site1}$, CENP-T$^{site2}$, CENP-T$^{\Delta N}$, CENP-T$^{flanks2/helix1}$ and CENP-T$^{flanks1/helix2}$ sequences, and GST-tag was removed in purified proteins.

The construct for mi3 particles was derived from the SpyCatcher-mi3-6xHis plasmid (Addgene plasmid #112255). A FLAG-tag sequence was inserted at the 3' end of mi3. The SNAP-SpyCatcher plasmid was derived from a plasmid encoding SNAP-GBP-6xHis (2). DNA fragment encoding GBP was substituted with the DNA encoding SpyCatcher. The Spy-tag-GBP-6xHis construct was created by replacing the SNAP encoding DNA sequence with the sequence for the Spy-tag AHIVMVDAYKPTK (2). The SunTag scaffolds were obtained from pcDNA4TO-mito-mCherry-24xGCN4_v1 (Addgene plasmid #60913). Scaffolds were cloned into lentiviral plasmids generated from Lenti-Cas9-2A-Blast (Addgene plasmid #73310). CENP-T$^{WT}$ for cell expression was obtained from pKG174 (1). CENP-T$^{T11A}$, CENP-T$^{T85A}$, and the scFv-sfGFP tag were synthesized by Twist Bioscience. The scFv-sfGFP-CENP-T constructs were cloned into a repair template for the AAVS1 “safe harbor” locus (pNM280).

#### Cell line generation

Cell lines were generated in a HeLa cell background using Cheeseman lab HeLa cells. Doxycycline-inducible scFv-sfGFP-CENP-T cell lines were generated by homology-directed insertion into the AAVS1 “safe-harbor” locus. Donor plasmid containing selection marker, the tetracycline-responsive promoter, the transgene, and reverse tetracycline-controlled transactivator flanked by AAVS1 homology arms (3) was transfected using Lipofectamine 2000 (Thermo Fisher Scientific) with a pX330-based plasmid (4) expressing both spCas9 and a guide RNA specific for the AAVS1 locus (pNM220; gRNA sequence – 5’-GGGGCCACTAGGGACAGGAT). Cells were selected with 0.5 µg ml$^{-1}$ puromycin (Life

Technologies). Clonal lines were obtained by fluorescence activated cell-sorting single cells into 96 well plates.

SunTag scaffolds were introduced by lentiviral transduction. Lentivirus was generated by using Xtremegene-9 (Roche) to co-transfect the scaffold-containing pLenti plasmid, VSV-G envelope plasmid, and Delta-VPR or psPAX2 (Addgene plasmid #12260) packaging plasmids into HEK-293T cells (5). Other lentivirus cell lines were selected with 2 µg $ml^{-1}$ blasticidin (Life Technologies). Cell lines containing SunTag scaffolds were generated from clonal parental lines expressing the desired sfGFP-scFv-CENP-T construct at comparable levels.

### Cell culture

HeLa cells were cultured in Dulbecco's modified Eagle medium (DMEM) supplemented with 10% fetal bovine serum (FBS), 100 U $ml^{-1}$ penicillin and streptomycin, and 2 mM L-glutamine at 37 °C with 5% $CO_2$. TetOn cell lines were cultured in FBS certified as tetracycline-free. TetOn constructs were induced with 1 µg $ml^{-1}$ doxycycline for 24 hours. To arrest cells in mitosis, cells were treated with 10 µM S-trityl-L-cysteine (STLC) for 16 hours. HeLa cells were regularly monitored for mycoplasma contamination.

### Immunofluorescence

Cells were seeded on poly-L-lysine (Sigma-Aldrich) coated coverslips. Cells were pre-extracted with 0.25% PBS-Tx (PBS with 0.25% Triton X-100), then fixed in 4% formaldehyde in PBS. Coverslips were washed with 0.1% PBS-Tx (PBS with 0.1% Triton X-100) and blocked in Abdil (20 mM Tris-HCl, 150 mM NaCl, 0.1% Triton X-100, 3% bovine serum albumin (BSA), 0.1% $NaN_3$, pH 7.5). Primary antibodies were diluted in Abdil. Centromeres were detected with anti-centromere antibodies (1:100 dilution; Antibodies, Inc, 15-234-0001), and Ndc80 complex was detected with anti-Bonsai antibodies (1:4,800 dilution; (6). Cy3- and Cy5-conjugated (or Alexa 647-conjugated) secondary antibodies (Jackson ImmunoResearch Laboratories) were diluted 1:300 in 0.1% PBS-Tx. DNA was stained with 1 µg ml Hoechst-33342 (Sigma-Aldrich) in 0.1% PBS-Tx. Coverslips were mounted with PPDM (0.5% *p*-phenylenediamine, 20 mM Tris-HCl, pH 8.8, 90% glycerol). Images were acquired with a DeltaVision Ultra High-Resolution microscope (Imsol). All images are maximal intensity projections in z. image manipulation was performed in Fiji (7).

Integrated fluorescence intensity of mitotic centromeres was measured with a custom CellProfiler 4.0 pipeline (8) (adapted from (9)). The median intensity of a 5-pixel region surrounding each centromere was multiplied by the area of the centromere to determine background intensity and subtracted from the integrated fluorescence of each centromere. Regions with high GFP signal were masked to avoid measuring kinetochore proteins bound to GFP-tagged constructs. Values for each cell were calculated from the mean of the Ndc80 complex signals of kinetochores in that cell. Before calculating the mean for a cell, the Ndc80 signal of each kinetochore in the cell was normalized to anti-centromere antibody signal from that kinetochore. Overall means and statistics were calculated from pooled data from multiple experiments. To make results comparable between experiments, the mean for each cell was normalized to the mean of all cells in the 1xGCN4pep sample in the same experiment. All image quantifications were performed on raw pixel values.

**DNA content analysis**

Cells were incubated in 1 μg ml$^{-1}$ doxycycline for 24 hours. 5 mM EDTA, 20 μg ml$^{-1}$ Hoechst-33342 (Sigma-Aldrich), and 10 μM Verapimil (Tocris; Spirochrome) were added directly to media for 30 minutes to 1 hour to detach cells from the plate and stain them. Cells were collected and filtered through 35 μm nylon mesh (Falcon). Hoechst, GFP, and tdTomato signals were measured on an LSRFortessa (BD Biosciences) flow cytometer. Results were analyzed with FlowJo software. The fraction of cells in each cell cycle phase was determined in FlowJo with a Watson (Pragmatic) model using the Cell Cycle tool. The DNA content of at least 5,000 cells was analyzed for each condition for each experiment.

**Protein expression and purification**

CENP-T constructs (CENP-T$^{6D}$, CENP-T$^{2D}$ and CENP-T$^{2D,short}$) were expressed in ArcticExpress (DE3) *Escherichia coli* (Agilent Technologies, 230192*)*. Expression was induced using 0.5 mM isopropyl β-D-1-thiogalactopyranoside (IPTG) and grown for 22 h at 10 °C. Cells were lysed by sonication in ice-cold Buffer A (50 mM Tris-HCl pH 8.0, 300 mM NaCl, 5 mM β-mercaptoethanol) supplemented with 0.15 mg ml$^{-1}$ lysozyme (Sigma-Aldrich, L6876), cOmplete EDTA-free protease inhibitors cocktail (Roche, 11873580001) and 1 mM phenylmethylsulfonyl fluoride (Sigma-Aldrich, P7626). Cell debris was cleared by ultracentrifugation in a Beckman 50.2Ti rotor at 50,000 *g* for 30 min, 4 °C. Cleared supernatant was applied to an equilibrated 5 ml HisTrap HP column (Cytiva, 17524801) on a fast protein liquid chromatography (FPLC) system AKTA Pure (GE Healthcare) at 4 °C. The column was washed with ten volumes of Buffer A, five volumes of the same buffer supplemented with 25 mM imidazole, and five volumes of buffer with 50 mM imidazole. The protein was eluted with ten column volumes of Buffer A containing the gradient of imidazole in the range 50-500 mM. Protein elution was monitored by absorbance at 280 nM and fractions containing CENP-T proteins were selected after sodium dodecyl sulfate-polyacrylamide gel electrophoresis (SDS-PAGE) analysis. Fractions containing CENP-T were combined, concentrated with 10 kDa Amicon Ultra Centrifugal Filter Units (EMD Millipore, UFC901008), and centrifuged to remove aggregates at 30,000 *g* for 15 min, 4 °C. The soluble fraction was applied to HiLoad Superdex 200 pg column (Cytiva, 28-9893-35) equilibrated with Buffer A on FPLC system AKTA Pure. Buffer A was applied at a flow rate 0.5 ml min$^{-1}$, all at 4 °C. Finally, purified proteins were supplemented with 20% glycerol, aliquoted, flash-frozen in liquid nitrogen, and stored at -80 °C.

CENP-T$^{site1}$, CENP-T$^{site2}$, CENP-T$^{\Delta N}$, CENP-T$^{flanks2/helix1}$and CENP-T$^{flanks1/helix2}$ constructs were expressed and purified following same procedure with several modifications. The lysis buffer was supplemented with cOmplete protease inhibitors cocktail (Roche, 11697498001) and 0.5 mM EDTA. The clear lysate was loaded onto a Glutathione Sepharose 4B column (Cytiva, 17-0756-01) that was pre-equilibrated with Buffer A containing 0.5 mM EDTA. The column was washed with 20 column volumes of Buffer A, and the proteins were eluted by cleavage from the column using 50 µg ml$^{-1}$ TEV protease in Buffer A with 0.5 mM EDTA overnight at 4 °C. TEV protease pTrc99 7xH TEV was a gift from Dr. Lampson, University of Pennsylvania. The fractions containing target proteins were concentrated and loaded onto a HiLoad Superdex 200 pg column. The SNAP-SpyCatcher, SNAP-GBP and Spy-tag-GBP proteins were expressed in BL21(DE3) *Escherichia coli* (NEB, C2527H). Expression was induced using 0.1 mM IPTG and grown for 18 h at 16 °C. Cells were lysed using a microfluidic chamber in ice-cold Buffer B (50 mM $NaH_2PO_4$ pH 8.0, 300 mM NaCl, 1 mM DTT, 0.1% Tween20 (Sigma-Aldrich, P1379) supplemented with 1 mg ml$^{-1}$ lysozyme, 1 mM 4-(2-aminoethyl)benzenesulfonyl fluoride hydrochloride (AEBSF, Santa Cruz Biotechnology, sc-202041A) and 10 mM imidazole.

After centrifugation at 50,000 g for 30 min, 4 °C supernatant was filtered and incubated with Ni-nitrilotriacetic acid (Ni-NTA) agarose (Qiagen, 30210) for 1 h at 4 °C. Bound protein was washed with Buffer B supplemented 20 mM imidazole and 1 mM AEBSF. The protein was eluted with Buffer B supplemented with 250 mM imidazole. To reduce the concentration of imidazole in the eluate, the buffer was changed to Buffer B without imidazole using Ultra Centrifugal Filter Units (EMD Millipore, UFC901008). Protein fractions were supplemented with 20% glycerol, aliquoted, snap-frozen in liquid nitrogen, and stored at -80 °C (*SI Appendix*, Fig. S1A).

SNAP-GBP was labeled with SNAP-Surface Alexa Fluor 647 (NEB, S9136S). The conjugation reaction was carried out in Buffer B without Tween20 using 12 µM Alexa Fluor 647 dye and 4 µM SNAP-GBP, with a 3-fold excess of dye over the protein. The reaction was allowed to proceed for 3 hours at room temperature. To separate unbound dye from labeled protein reaction mixture was loaded to PD-10 column (Cytiva, 17085101) column and eluted by Buffer B without Twin20. Finally, SNAP-GBP labeled with Alexa Fluor 647 was supplemented with 20% glycerol, aliquoted, flash-frozen in liquid nitrogen, and stored at -80 °C. In the paper for simplicity we call this protein “GBP-Alexa Fluor647”. The degree of labeling was estimated as the ratio of the Alexa Fluor 647 concentration (measured at 647 nm) to the protein concentration (measured at 280 nm), yielding 0.8.

Human Ndc80 Bonsai complex containing the N-terminal fragment of Hec1 (1-286 aa) fused to a fragment of the Spc25 (118-224 aa) with C-terminal GFP or untagged and the Nuf2 protein (1-169 aa) fused to a fragment of Spc24 (122-197 aa) was expressed in *Escherichia coli* and purified, as in (10) (*SI Appendix*, Fig. S1A). Human Ndc80 ΔSpc24/25 complex containing N-terminal fragments of Hec1 (1-506 aa) and Nuf2 (1-348 aa) with GFP-tag on the C-terminus of Nuf2 was expressed and purified, as in (6).

The protocol for purification of mi3 core particles (6xHis-SpyCatcher-mi3) was based on (11). Mi3 core particles were expressed in BL21(DE3) *Escherichia coli*. Expression was induced using 0.5 mM IPTG overnight at 22 °C. Cells were lysed and sonicated in ice-cold Buffer C (50 mM Tris-HCl pH 8.5, 500 mM NaCl) supplemented with 0.15 mg $ml^{-1}$ lysozyme, cOmplete EDTA-free protease inhibitors cocktail and 1 mM phenylmethylsulfonyl fluoride. Cell debris were cleared by ultracentrifugation in a Beckman 50.2Ti rotor at 50,000 *g* for 30 min, 4 °C. Cleared supernatant was applied to equilibrated 5 ml HisTrap HP column on a FPLC system AKTA Pure at 4 °C. The column was washed with ten volumes of Buffer C, five volumes of the same buffer supplemented with 25 mM imidazole, and five volumes of buffer with 50 mM imidazole. The protein was eluted with ten column volumes of Buffer C containing the gradient of imidazole in range of 50-500 mM. Protein in elution was monitored by absorbance at 280 nM and fractions containing 6xHis-SpyCatcher-mi3 were selected after SDS-PAGE-analysis. Fractions containing SpyCatcher-mi3 were combined, concentrated to 5 ml with 100 kDa Amicon Ultra Centrifugal Filter Units and centrifuged to remove aggregates at 30,000 *g* for 15 min, 4 °C. The soluble fraction was applied to HiPrep 16/60 Sephacryl S-400 column equilibrated with Buffer C on FPLC system AKTA Pure. Buffer C was applied at flow-rate 0.5 ml $min^{-1}$, all at 4 °C. Based on the SDS-PAGE-analysis, the fractions containing 6xHis-SpyCatcher-mi3 particles are in the 60-75 ml range. Mi3-based core particles were supplemented with 20% glycerol, aliquoted, flash-frozen in liquid nitrogen and stored at -80 °C.

**TIRF microscopy assays to study interactions between monomeric CENP-T and soluble Ndc80**

Flow chamber preparation and imaging. Prior to each experiment, frozen protein aliquots of SNAP-SpyCatcher, Ndc80, and CENP-T were thawed on ice and clarified by ultracentrifugation at 156,000 *g* for 15 min at 4 °C. Flow chambers were prepared as described in (12) using silanized coverslips and reusable glass slides with tubing (13). To perfuse solutions into the flow chamber, a syringe pump (New Era Pump Systems, NE-4000) was used to draw the liquid at a speed of 150 μL $min^{-1}$, unless otherwise indicated. The specimen on the microscope stage was maintained at 32 °C. The coverslip of the assembled and sealed flow chamber was functionalized by incubation with 100 nM of SNAP-SpyCatcher in PBS buffer (10.1 mM $Na_2HPO_4$, 1.8 mM $KH_2PO_4$ pH 7.2, 140 mM NaCl 2.7 mM KCl) supplemented with 2 mM DTT for 10 min. Next, the chamber surface was blocked with 1% Pluronic F127 (Sigma-Aldrich, CP2443). Then, 1-3 nM of CENP-T-GFP-Spy-tag in PBS buffer supplemented with 2 mM DTT, 4 mg $ml^{-1}$ BSA (Sigma-Aldrich, A7638) and 0.5 mg $ml^{-1}$ casein (Sigma-Aldrich, C5890) were introduced for 5 min to achieve the desired density of CENP-T molecules on the coverslip (Fig. 1C). Chambers were then incubated with Imaging Buffer: Mg-BRB80 (80 mM K-PIPES pH 6.9, 1 mM EGTA, 4 mM $MgCl_2$) supplemented with 10 mM DTT, 4 mg $ml^{-1}$ BSA and 0.5 mg $ml^{-1}$ casein, 0.1 mg $ml^{-1}$ glucose oxidase (Sigma-Aldrich, G2133), 20 μg $ml^{-1}$ catalase (Sigma-Aldrich, C40) and 6 mg $ml^{-1}$ glucose (Sigma-Aldrich, G8270).

All fluorescent imaging was performed using a Nikon Eclipse Ti microscope equipped with a 1.49xNA TIRF 100x Oil objective. Excitation for visualizing GFP- and Alexa Fluor 647-tagged proteins in TIRF modes was provided by coherent CUBE 488-nm and 640-nm 50 mW diode lasers (Coherent). Images were captured using an Andor iXon3 EMCCD camera. The concentration of GFP-tagged proteins was determined by measuring GFP intensity through fluorescence microscopy, as described in (12).

Before the addition of Ndc80, a set of images of immobilized GFP-tagged CENP-T molecules in several imaging fields was captured to determine their initial coordinates and intensities. Subsequently, 200 μl of 200 nM GFP-tagged Ndc80 in Imaging Buffer were introduced using a syringe pump at a speed of 900 μl $min^{-1}$. The same fluorophore was used on CENP-T and GFP molecules to simplify intensity quantification and avoid errors caused by uneven or varying illumination. The interaction time between Ndc80 and CENP-T varied in experiments, ranging from 2 to 60 minutes. At specific time points, a second image of one of the initial fields was captured to observe the brightness of the dots. At indicated times, soluble Ndc80 was washed out by perfusing 200 μl of Imaging Buffer at a speed of 900 μl $min^{-1}$. Then, the pump speed was reduced to 5 μl $min^{-1}$ and the flow was maintained to remove any remaining Ndc80 molecules that detached from CENP-T clusters over time. During the unbinding phase, images of a different set of initially captured fields were collected at the desired time points.

Data analysis. The images of CENP-T molecules were analyzed using the Fiji software (7). To confirm that immobilized CENP-T molecules are monomeric, the distribution of their initial fluorescence intensities was determined. Specifically, fluorescence intensity was measured in the area surrounding the CENP-T molecule (3-pixel radius). The brightness of the same-sized area located near the selected molecule was subtracted to minimize variability in background intensity. The fluorescence intensities of individual GFP fluorophores were determined using photobleaching curves (*SI Appendix*, Fig. S1B,C), as described in (14). The number of GFP molecules per CENP-T dot was determined as a ratio of CENP-T intensity dot and intensity of one GFP molecule (*SI Appendix*, Fig. S1E-H). Analysis of GBP-Alexa Fluor 647 was done analogously (*SI Appendix*, Fig. S1D).

To quantify Ndc80 binding to the coverslip-immobilized CENP-T monomers, the pair of images before and after Ndc80 addition was manually corrected for a stage drift. Then, the 30-40 GFP-tagged CENP-T molecules were manually selected on the initial CENP-T image. If a CENP-T molecule detached during the experiment, it was not considered for further analysis. The number of bound Ndc80 molecules was calculated as the difference between the intensity after Ndc80 binding and the initial CENP-T-GFP intensity, divided by the initial intensity. For each time point corresponding to the pair of experimental images, the median Ndc80 binding was determined. Next, the median Ndc80 binding was plotted against time, and specific portions of the curve representing Ndc80 binding and unbinding were fitted using one-component exponents (Fig. 1D,E). For the unbinding exponent, the initial point was set to be similar to the plateau of the binding exponent. The total binding level was determined as the plateau of the exponent fitting the binding curve, while the stable binding level was determined as the plateau of the exponent fitting the unbinding curve. The $k_{off}$ rate was obtained as the exponent parameter of the unbinding curve fitting. The transient binding was determined as a difference between total and stable binding.

To evaluate the binding of GBP-Alexa Fluor647 to GFP-tagged CENP-T the analogues procedure was done with the following modification. First, the number of CENP-T molecules for one GFP dot was quantified by dividing the dot's fluorescence intensity by the intensity of one GFP molecule. Then, the number of GBP-Alexa Fluor647 molecules was quantified by dividing the dot's fluorescence in the second fluorescent channel by the intensity of one Alexa Fluor647 molecule. Finally, the number of GBP-Alexa Fluor647 molecules was divided by the number of CENP-T molecules to get the efficiency of binding.

Estimation of photobleaching effect. Since photobleaching of GFP molecules on CENP-T can potentially affect the accurate quantification of Ndc80 binding to CENP-T, imaging conditions were selected to minimize the probability of photobleaching. The illumination time was minimized to 0.3 s, and each experimental field was captured only twice: initially before Ndc80 addition and at a specific time after Ndc80 addition. To estimate probability of photobleaching of GFP molecules over this illumination time (0.6 s), the rate of photobleaching was measured using GFP-tagged CENP-T$^{6D}$ molecules immobilized on the coverslip. Imaging field with GFP-tagged CENP-T$^{6D}$ molecules was continuously illuminated using the same illumination settings as those used in experiments with CENP-T monomers. The number of GFP dots per imaging field at each time point was counted and normalized to the number of dots in the first imaging frame. Data from N = 3 independent experiments were combined; the fraction of fluorescent molecules was calculated as a number of fluorescent dots at the indicated time point normalized on an initial number of fluorescent dots. The resultant curve was fitted to an exponential decay function to determine the characteristic photobleaching time (*SI Appendix*, Fig. S2B, left panel). The probability of photobleaching was then calculated as the illumination time 0.6 s multiplied by the exponent coefficient, resulting in 6%. This probability is within the range of experimental error from multiple experimental repeats, so it is not expected to significantly affect the results.

**TIRF microscopy assay to study interactions between clustered CENP-T and soluble Ndc80**

Preparation of CENP-T clusters in a flow chamber and imaging. The flow chamber was constructed as in experiments with CENP-T monomers. After the chamber was blocked with Pluronic F127, mi3 particles (100 nM) were allowed to adsorb on the coverslip for 10 min. Next, 200 nM of CENP-T in PBS buffer supplemented with 2 mM DTT, 4 mg $ml^{-1}$ BSA and 0.5 mg $ml^{-1}$ casein were introduced for 20 min at room temperature. Chamber with assembled CENP-T clusters was rinsed with Imaging Buffer.

Control Ndc80-containing clusters were assembled analogously. First, 75 nM Spy-tag-GBP in PBS buffer supplemented with 2 mM DTT, 4 mg $ml^{-1}$ BSA, and 0.5 mg $ml^{-1}$ casein was incubated with adsorbed mi3 particles for 15 min at room temperature. After washing, 500 nM GFP-tagged Ndc80 Bonsai was incubated for 30 min in the same buffer, the chamber was washed with Imaging Buffer.

Images of clusters were acquired in TIRF mode with a 488 nm 100 mW diode laser (Coherent, Santa Clara, CA, USA) at 1% power with an exposure time of 30 ms. After the GFP-tagged CENP-T clusters were assembled, several images were collected for subsequent quantifications of initial fluorescence intensity, corresponding to the quantity of CENP-T molecules per cluster. The clusters were then bleached with laser at 100% power for 30 s. Binding and unbinding of Ndc80 was imaged as 2-60 images per minute, with less frequent image capture employed during longer incubations to reduce photobleaching. Unlike single molecule settings, the laser power was significantly reduced for cluster imaging, and there was no requirement to capture different fields for different time points. Some experiments with CENP-T clusters were carried out in the presence of a continuous flow (30 µl $min^{-1}$) of Imaging Buffer with Ndc80 protein, but in others the flow was used only to introduce Ndc80 into the chamber. This variation did not affect final binding results, so the data for stable Ndc80 binding were combined.

<u>Data analysis.</u> To evaluate level of Ndc80 binding, image sequences with CENP-T clusters were analyzed similarly to monomers with several modifications. First, for stage drift correction of the image stack the “Manual drift correction” plugin in Fiji software was used (7). Second, the area in which fluorescence intensity was measured was increased up to 8-pixel radius to fit CENP-T clusters. Finally, the initial fluorescence intensity of clusters was not subtracted from intensity of clusters after Ndc80 addition due to bleaching of GFP molecules on CENP-T clusters. Typically, approximately 30 clusters were analyzed for each independent experiment.

<u>Correction for photobleaching.</u> To correct for GFP-photobleaching, the rate of photobleaching for GFP-tagged CENP-T clusters was determined. GFP-tagged CENP-$T^{6D}$ clusters immobilized on the coverslip were continuously illuminated and imaged with microscope settings identical to those used in experiments with CENP-T clusters. Integral intensity of clusters was determined for each time point and normalized on the initial cluster intensity. The curve for clusters’ intensity vs. illumination time was fitted with exponential decay function to generate the control photobleaching curve (*SI Appendix*, Fig. S5C). To take into account photobleaching of GFP during experiments with Ndc80 and CENP-T, all curves corresponding to Ndc80 unbinding during the washout were normalized on the control photobleaching curve. During the biding phase, soluble Ndc80-GFP molecules continuously exchange with CENP-T, so the intensities collected in the presence of soluble Ndc80-GFP were not corrected for photobleaching. Total illumination time during the binding phase was < 3.6 s, so bleaching would affect < 5% of GFP-containing molecules.

<u>Quantification of the size of CENP-T clusters.</u> The size of GFP-tagged CENP-T clusters was calculated based on their fluorescence intensity. First, the intensity of individual GFP-molecule was captured using the following settings for Andor iXon3 camera: 10 MHz readout speed, gain 5.0x, EM gain 999, 30 msec exposure time; and 50% power of 488 nm 100 mW diode laser (Coherent, Santa Clara, CA, USA), as described in (14). Next,

images of the GFP-tagged CENP-T clusters were captured using the same camera settings, except the laser power which was reduced from 50% to 1% to avoid camera saturation. Then, the images of clusters were corrected using the laser intensity profile, which was obtained, as described in (14), by averaging images with high density of CENP-T clusters on the coverslip. Finally, number of GFP-tagged CENP-T molecules per cluster was calculated as ratio of fluorescence intensity of individual clusters divided by the intensity of single GFP molecule and multiplied by 31.2 which resulted from different laser power settings (*SI Appendix,* Fig. S5B).

Analysis of Ndc80 binding to clusters containing Ndc80. To assess the extent of Ndc80-Ndc80 binding in our experiments with coverslip immobilized clusters we generated clusters containing Ndc80 with no CENP-T. These clusters were formed by fusing the linker protein Spy-tag-GBP to SpyCatcher-mi3 clusters, and then incubating with GFP-Ndc80 Bonsai for GFP/GBP binding. After washing out the unbound proteins, these Ndc80 clusters were tested using the same protocol as with the CENP-T clusters.

### FRAP assay to study interactions between clustered CENP-T and soluble Ndc80

CENP-$T^{6D}$ clusters were assembled on the coverslip, imaged, and photobleached, as during a regular TIRF-based assay. GFP-Ndc80 Bonsai (200 nM) was incubated for 2 or 60 minutes, and 25 image frames were collected to determined total Ndc80 binding in the presence of soluble Ndc80, which was 1.88 ± 0.02 Ndc80 molecules per CENP-T for both conditions (Fig. 2F). Imaging field was then photobleached with a 488-nm laser at 100% power for 45 seconds. The time between photobleaching and the start of imaging was 7 seconds. The recovery curves were fitted with an exponential function. The exchangeable fraction was determined as the plateau of this fitting, whereas the stable binding was calculated as the difference from total binding and the mobile fraction.

### FCS assay to study interactions between soluble Ndc80 and CENP-T proteins

FCS measurements were performed on a multi-parameter time-resolved confocal microscopy and spectroscopy instrument (MicroTime-200, PicoQuant, Germany). GFP-tagged CENP-T proteins were excited with 484 nm pulsed diode laser (LDH-D-TA-484, PicoQuant) operating at 20 MHz, through an excitation dichroic filter ZT440-445/484-491/594 rpc-UF3 (Chroma Technology) and an Olympus UPLanSApo 60x/1.2-W water objective lens. The laser power was maintained at ~15 µW and it was focused 20 µm above the coverslip interface for measurement. After passing through a 50 µm pinhole, the fluorescence signals from the sample were split into two channels by a polarized beam splitter (U-MBF3-Olympus). The signals further passed through bandpass filters ET535/70 (Chroma Technology) and projected onto two single-photon avalanche photodiode detectors (SPAD: SPCM-AQRH-14-TR). Nunc Lab-Tek chambers (ThermoFisher, 155411) with borosilicate cover slip bottoms were used at 19-21 °C. The chambers were passivated by treatment with 50% PEG-8000 solution, incubated at room temperature for 3-4 hours, followed by 3-4 washes with the Mg-BRB80 buffer. Atto-488 dye was used as a standard sample to calibrate the confocal detection volume of the 484 nm laser beam.

Ndc80 binding experiments were carried out by rapid mixture of 1 nM GFP-tagged CENP-T with 0-1000 nM of unlabeled Ndc80 Bonsai in Mg-BRB80 buffer supplemented with 2 mM DTT and 4 mg $ml^{-1}$ BSA and 0.5 mg $ml^{-1}$ casein.

The analysis of time-trace curves was performed using the SymPhoTime software provided by PicoQuant. To ensure accurate results, the time-trace signals were filtered with an 80-count threshold to eliminate aggregates, followed by background correction (*SI*

*Appendix*, Fig. S3A). Cross-correlation curves were generated using signals from two detectors and fitted with a triplet model (*SI Appendix*, Note 1.3, Fig. S3B-D) (15).

#### Negative staining electron microscopy

For electron microscopy (EM) experiments CENP-$T^{2Dshort}$, clusters were assembled in a test tube and subsequently purified using gel-filtration chromatography (*SI Appendix*, Fig. S4A). Briefly, 80 nM of SpyCatcher-mi3 core particles were conjugated to 10 µM of Spy-tagged CENP-$T^{2D,short}$. The conjugation reaction was carried out in buffer D (50 mM Tris-HCl pH 8.0, 150 mM NaCl) for 3 hours at room temperature. Separation of the conjugated clusters from unconjugated CENP-T was performed by loading the reaction mixture onto a HiPrep 16/60 Sephacryl S-400 column. The column was equilibrated and washed with buffer D at a flow rate of 0.5 ml $min^{-1}$. The CENP-$T^{2D,short}$ clusters were eluted in the range of 60-75 ml, supplemented with 20% glycerol, aliquoted, flash-frozen in liquid nitrogen, and stored at -80 °C. For imaging, CENP-$T^{2D,short}$ clusters were applied to freshly glow-discharged carbon-coated 200 mesh copper grids (Electron Microscopy Sciences, CF200-CU-50) and incubated for 1 minute. Excess liquid was blotted off using filter paper. The grids were washed three times and stained with 2% uranyl acetate for 30 seconds. After staining, the excess stain was blotted off, and the grids were air-dried. Imaging of the grids was performed using a JEM 1011 transmission electron microscope (JEOL) operated at 100 kV, coupled with an ORIUS 832.10W CCD camera (Gatan). Clusters with no conjugated proteins were used as control. The size of clusters was estimated with Fiji software (*SI Appendix*, Fig. S4B).

#### Mathematical modeling

Theoretical approaches and our model are described in *SI Appendix*, Note 2. Briefly, interactions between Ndc80 and CENP-T were analyzed by determining numerical solutions of a system of differential equations for different proteins and initial conditions. The model predicted fraction of Ndc80/CENP-T complexes as a function of time and Ndc80 concertation. MATLAB version R2020b with a MATLAB ODE solver *ode45* were used.

#### Structural analysis of Ndc80/CENP-T complexes using AlphaFold software

Analysis of chicken proteins and human Spc24/Spc2 were conducted using AlphaFold2.2.4 (16). Simulations were performed using the 'alphafold2_multimer_v3' model, recycling five models until the predicted structural tolerance fell below 0.5, after which Amber relaxation (17) was applied to the highest-ranked model. Structural predictions for human Ndc80 in a complex with CENP-T binding sites, as well as CENP-T binding sites with no NDc80, were generated using AlphaFold3 via the AlphaFold3 server (18). Input sequences were from the PDB database: 3VZA for chicken Spc24(134-195 aa)/Spc25(134-232 aa) in a complex with CENP-T site (63-93 aa; T72D, S88D) (19), and 2VE7 for human Spc24(122-197 aa)/Spc25(118-224 aa) (20). Human CENP-T sequences were 1-30 aa for site 1 and 76-106 aa for site 2 (NCBI: KAI2579221.1); both sequences were modified with phosphomimetic substitutions (T11D and T85D), as used in our experiments). The analysis and alignments of the predicted structures were conducted using the PyMOL Molecular Graphics System, Version 2.5 (Schrödinger, LLC). To quantify structural similarities, the template modeling (TM) score was calculated using the Python tmtools package version 0.0.3 (21, 22). To estimate reproducibility of model predictions, five independent simulations for each model were carried out and the standard deviation (SD) of the Cα atoms coordinates of each residue in CENP-T was calculated using a Python script based on Pandas data analysis library and Biopython. For results and discussion see *SI Appendix*, Note 3.

### Supplementary text

#### Note 1. Supporting experimental results

1.1 CENP-T does not mature on its own. Experiments using coverslip-immobilized CENP-T revealed a time-dependent increase in the fraction of stably bound Ndc80 molecules. One possible explanation for this increase is that prolonged incubation of CENP-T on the coverslip surface could modify its properties, altering its affinity for Ndc80. To test this, we incubated immobilized CENP-$T^{6D}$ with imaging buffer for 60 minutes, then added 200 nM Ndc80. After 2 minutes, the Ndc80 was washed out, and both the total and stable Ndc80 binding to CENP-$T^{6D}$ were measured. The results were the same as when Ndc80 was added immediately after immobilization (*SI Appendix*, Fig. S2A). Thus, any slow changes in CENP-T's ability to retain Ndc80 (i.e., maturation) are not due to incubation alone and require continuous interactions with Ndc80. This supports the two-step reaction mechanism for Ndc80/CENP-T complex formation.

1.2 Stability of Ndc80 complex with CENP-T monomers in the presence of soluble Ndc80.

To assess the stability of Ndc80/CENP-T complexes in the presence of soluble Ndc80, CENP-$T^{6D}$ monomers were immobilized on coverslips using a standard TIRF-based microscopy assay. GFP-Ndc80 Bonsai was added at 200 nM, followed by incubation for either 2 or 60 minutes. GFP fluorescence imaging was then performed continuously for 5 minutes under illumination conditions that induce GFP photobleaching with a characteristic photobleaching time of 10 seconds (*SI Appendix*, Fig. S2B, left panel). In control experiments without soluble GFP-Ndc80, fluorescence from immobilized CENP-T quickly faded. However, in the presence of soluble Ndc80, the fluorescent signal persisted much longer, indicating binding of unbleached GFP-Ndc80 to the immobilized CENP-T monomers (*SI Appendix*, Fig. S2B, kymographs and intensity profiles). No GFP fluorescence was detected when soluble GFP-Ndc80 was present without CENP-T, confirming that the signal arose from Ndc80 binding to CENP-T. The observed blinking of fluorescence indicated repeated cycles of GFP-Ndc80 binding to unoccupied CENP-T sites, followed by Ndc80 unbinding or photobleaching. Interestingly, the intensity of blinking was higher in samples incubated for 2 minutes, but its frequency decreased after 60 minutes of interaction. These results strongly suggest a time-dependent change in the exchange rate between Ndc80 and CENP-T, confirming our conclusions from experiments that removed soluble protein. The Ndc80 washout approach provided a more precise measurement of Ndc80 binding and unbinding dynamics, making it the preferred method for our quantitative study.

1.3 Interactions between Ndc80 and CENP-T in solution analyzed with FCS

To further test the conclusions from our experiments using the TIRF-based assay with immobilized CENP-T monomers, we turned to the Ndc80/CENP-T binding kinetics in solution, where molecular interactions are free from possible interference from surface proximity. Fluorescence Correlation Spectroscopy (FCS) (23) was used to determine binding kinetics between soluble unlabeled Ndc80 Bonsai and GFP-tagged CENP-$T^{2D}$, as well as CENP-T mutants lacking one of the sites.

First, we determined the diffusion time for GFP-tagged CENP-T molecules (CENP-$T^{2D}$, CENP-$T^{site1}$ or CENP-$T^{site2}$) using these proteins at 1 nM. *SI Appendix*, Fig. S3, panels B-D show the corresponding cross-correlation curves. Since GFP-fluorophore undergoes transition into a triplet state (24), these curves were fitted with the one-component triplet models. The diffusion time, which was a fitting parameter (25) was thus determined for each protein: 0.42 ± 0.03 ms for CENP-$T^{2D}$, 0.41 ± 0.02 ms for CENP-$T^{site1}$, 0.38 ± 0.02 ms for CENP-$T^{site2}$ (*SI Appendix*, Fig. S3E). These values are similar, consistent with the similar molecular size of these CENP-T proteins.

Next, we determined the diffusion time for Ndc80/CENP-T complexes using GFP-tagged CENP-T proteins and unlabeled Ndc80 Bonsai protein. To ensure that most CENP-T molecules in the reaction mixtures are bound to Ndc80, we used high concentration of Ndc80. For CENP-$T^{2D}$ we used 20 nM and 100 nM Ndc80; for CENP-$T^{site1}$ – 100, 300 and 1,000 nM; for CENP-$T^{site2}$ – 100 nM. The cross-correlation curves for the reaction mixtures containing a CENP-T protein with different Ndc80 concentrations were highly similar, indicating saturation of the CENP-T binding sites. These curves were shifted relative to signals from the corresponding CENP-T proteins with no Ndc80 (*SI Appendix*, Fig. S3B-D), indicating increase in the diffusion times due to the larger sizes of the Ndc80/CENP-T complex vs. CENP-T alone. These cross-correlation curves were then fitted using a one-component triplet models. The resultant diffusion times for different Ndc80 concentrations were similar for each CENP-T protein, as expected. These values were averaged to represent the diffusion time for each type of complexes (*SI Appendix*, Fig. S3E): 0.61 ± 0.03 ms for CENP-$T^{2D}$ in complex with two Ndc80s, 0.49 ± 0.04 ms for Ndc80/CENP-$T^{site1}$, 0.47 ± 0.02 ms for Ndc80/CENP-$T^{site2}$. For all CENP-T proteins, the binding of Ndc80 significantly slowed down the rate of CENP-T diffusion (higher diffusion time) without significantly changing complex intensity (*SI Appendix*, Fig. S3E,F), consistent with binding of non-fluorescent Ndc80 molecules and presence of only one copy of CENP-T in the assembled complexes, as expected.

Using these diffusion time values for CENP-T proteins alone and in a complex with Ndc80, we then determined the kinetics of Ndc80 binding to CENP-T. Mixtures of 1 nM GFP-tagged CENP-T and 10 nM unlabeled Ndc80 were prepared and FCS data were collected at 5, 10, 30, 45, 60 and 90 min. To analyze reaction mixtures containing the CENP-T proteins with only one Ndc80 binding site (CENP-$T^{site1}$ and CENP-$T^{site2}$) we applied a two-component triplet model using as fixed parameters the diffusion time values for CENP-T monomers and the corresponding Ndc80/CENP-T complexes. The model output was the fraction of Ndc80/CENP-T complexes in the reaction mixture as a function of time. Since CENP-$T^{2D}$ protein has two Ndc80 binding sites, the cross-correlation curves for the reactions with this protein were analyzed using a three-component triplet model. The diffusion times for CENP-$T^{2D}$ alone and in a complex with two Ndc80 molecules were fixed using their independently determined values, as explained above. The diffusion time for complexes containing one Ndc80 molecule (in site 1 or site 2) were unknown. For simplicity, we assumed that these two complexes have the same diffusion time, and that it is similar to the diffusion time measured for the complexes with the mutant CENP-T proteins (*SI Appendix*, Fig. S3G). We averaged the experimentally determined diffusion time values for complexes between Ndc80 and CENP-$T^{site1}$ or CENP-$T^{site2}$, and used this value as a third parameter for the three-component triplet model. This model failed to converge, likely because of the relatively minor difference between the diffusion time for CENP-$T^{2D}$ and the diffusion times for complexes containing Ndc80 and CENP-T mutants with only one site (*SI Appendix*, Fig. S3G). To overcome this problem, we simplified the fitting model to include only two components, corresponding to the slow and fast diffusing molecular species. The slow diffusing component is produced by the largest molecular complex: CENP-$T^{2D}$ bound to two Ndc80 molecules (diffusion time 0.61 ± 0.03 ms). The fast-diffusing component corresponds to a mixture of several proteins with similar diffusion times: CENP-$T^{2D}$ alone, CENP-$T^{2D}$ bound to one Ndc80 at either site 1 or site 2. The diffusion time for this component was obtained by averaging the values obtained for

CENP-T$^{2D}$ alone and for Ndc80/CENP-T$^{site1}$ and Ndc80/CENP-T$^{site2}$ complexes: 0.46 ± 0.02 ms. This model provided reliable and reproducible fitting, allowing us to estimate the fraction of CENP-T$^{2D}$ in complex with two Ndc80s as a function of reaction time.

To determine concentration dependencies for reactions involving GFP-tagged CENP-T molecules (CENP-T$^{2D}$, CENP-T$^{site1}$ or CENP-T$^{site2}$), the mixtures containing 1 nM CENP-T and 0-300 nM Ndc80 were prepared, and samples were collected for FCS analysis after 70 min interaction. The fraction of complexes was determined using the corresponding models, as described above. The concentration dependencies, as well as the kinetic curves, matched well the predictions of the model with maturing sites for the corresponding proteins (Fig. 1J,K). Thus, the results from two independent fluorescence-based assays (TIRF and FCS) support the proposed model of interaction between Ndc80 and CENP-T with binding site maturation.

### Note 2. Mathematical model of Ndc80 interaction with CENP-T

#### General overview of theoretical approaches

In our experiments, dissociation of Ndc80/CENP-T complexes is described by biphasic kinetics, implying the presence of two types of complexes with distinct dissociation constants. As the fraction of stable complexes increases over time, the initial complexes (referred to in the model as "nascent" binding sites) appear to transform into more stable complexes (with "mature" sites) through a stochastic site transition. We formalized these findings using the following reaction scheme for an Ndc80 binding site:

$$N + T \underset{k_{off}}{\overset{k_{on}}{\rightleftarrows}} (NT) \overset{k_{mat}}{\rightarrow} (NT)^* \underset{k^*_{off}}{\rightarrow} N + T \qquad \text{(S1)}$$

where $N$ corresponds to Ndc80, $T$ to Ndc80 binding site on CENP-T, $(NT)$ and $(NT)^*$ to intermediate and mature Ndc80/CENP-T complexes, correspondingly. Rate constants $k_{on}$ and $k_{off}$ describe Ndc80 association and dissociation with the nascent CENP-T site, $k^*_{off}$ describes the Ndc80 dissociation from mature Ndc80/CENP-T complex and $k_{mat}$ describes the transition from nascent to mature Ndc80/CENP-T complex.

This scheme is applicable to any of the two Ndc80 binding sites on CENP-T, but for the molecule with both sites the reaction transition scheme is more complex (*SI Appendix*, Fig. S7A). We constructed a chemical kinetics model that includes both sites and their respective transitions. To enable comparison of model predictions with our experimental results, the model considers interactions between soluble Ndc80 protein and CENP-T molecules immobilized on a coverslip surface. The applicability of the mass action law to surface-immobilized molecules and our use of probability-based modeling in conjunction with the fractions of interacting molecules are addressed below. The model's output is presented as the number of Ndc80 molecules bound to a single CENP-T molecule. This allows the modeling results to be directly compared with experimental kinetic curves obtained for coverslip-immobilized CENP-T monomers and clusters without additional model modifications. The model makes no assumptions about the nature of Ndc80 binding to monomers versus clusters of CENP-T, assuming identical binding mechanisms. All kinetic constants in the model were determined by fitting the results to experimental kinetic curves generated in experiments with coverslip-immobilized CENP-T clusters containing only one of the CENP-T sites (site1 and site2 proteins) (see section 'Model calibration').

To predict reactions between Ndc80 and CENP-T in solution the model equations more modified accordingly, as described in the section 'Model for soluble Ndc80 and CENP-T proteins'.

#### Application of the mass action law to reactions with the surface-immobilized molecules

According to the mass action law, the rate of a chemical reaction is directly proportional to the concentrations of the reactants. This law is well-suited for systems with a large number of reacting particles in a homogeneous medium. However, this approach needs modification when studying reactions involving surface-associated molecules. To adjust the mass action equation for a surface-bound reactant $A$, we analyzed the likelihood of interaction between molecule $A$ and a freely diffusing molecule $B$ in the solution. Similar to reactions occurring in solution, we assumed that the interaction between $A$ and $B$ occurs when the two molecules collide and subsequently undergo a chemical reaction. Consequently, the probability of interaction between molecules $A$ and $B$ (denoted as $p^{inter}$) is determined by the product of probabilities of their collision $p^{AB}$ and the subsequent chemical reaction $p^{react}$.

To calculate the probability of a collision, we first considered a general scenario in which molecules of reactant $A$ are distributed across a surface area $S$ exposed to a homogeneous solution of the freely floating molecules $B$ (*SI Appendix*, Fig. S7B). Assuming that the surface distribution of molecules $A$ is random, and the distance between the adjacent molecules is larger than the radius of effective interaction area $S^{eff}$, the probability $p^{smB}$ of one molecule $B$ with diameter $d_B$ to collide with the immobilized molecule $A$ with diameter $d_A$ is given by:

$$p^{smB} = \frac{S^{eff}}{S} = \frac{\frac{1}{4}\pi(d_A+d_B)^2}{S} \qquad \text{(S2)}$$

The probability of collision $p^{AB}$ between any soluble molecule $B$ and any immobilized molecule $A$ should be proportional to the concentration of molecules $B$, denoted by $[B]$. We assume that $[B]$ is sufficiently high to neglect any changes due to complex formation with molecules A, so $[B] = [B_0]$, where $[B_0]$ is the initial concentration of molecules $B$. Additionally, $p^{AB}$ should be proportional to the surface density of molecules $A$ that have not yet formed a complex with $B$, denoted by $< A >$. Thus, the probability of collision $p^{AB}$ is given by:

$$p^{AB} = p^{smB} < A > [B_0] \qquad \text{(S3)}$$

$$p^{AB} = \frac{\frac{1}{4}\pi(d_A+d_B)^2}{S} < A > [B_0] \qquad \text{(S4)}$$

Thus,

$$p^{inter} = p^{react}p^{AB} \sim < A > [B_0] \qquad \text{(S5)}$$

Direct application of equation (S5) to experiments that monitor binding between immobilized and soluble molecules is challenging. Indeed, the results of such experiments are not represented by the reactants concentrations, and the initial surface density of molecules $A$, denoted by $< A_0 >$, is often unknown. Moreover, when studying single molecules, the binding outcome is binary, indicating whether the complex has formed or not, further complicating the application of equation (S5). To address these challenges, we considered the fraction of immobilized single molecules $A$ that have not formed a complex with molecule $B$, denoted by $\{A\}$. We used the normalized probability of interaction $p^{measured}$, which is independent of the initial density $< A_0 >$:

$$p^{measured} = \frac{p^{inter}}{<A_0>} \sim \frac{<A>[B_0]}{<A_0>} \sim \{A\}[B_0] \qquad \text{(S6)}$$

Thus, the reaction rate $v$, which is proportional to $p^{measured}$, is given by

$$v = k\, p^{measured} \sim k\, \{A\}[B_0], \qquad \text{(S7)}$$

where $k$ is the corresponding kinetic constant.

Equation (S7) enables the construction of a system of differential equations to describe interactions between the coverslip-immobilized CENP-T molecules and soluble Ndc80 molecules. Moreover, the solutions of this system can be directly compared to experimental data for experiments using CENP-T monomers and clusters.

### Model for soluble Ndc80 and coverslip-immobilized CENP-T monomers and clusters

#### *Model framework*

Ndc80 protein. In the model, the concentration of soluble Ndc80 protein (in nM), denoted by $[N]$ was constant and equal to its initial concentration $[N_0]$, which was selected according to experimental conditions. Here and below the square brackets stand for the concentration of the corresponding soluble reactant.

CENP-T protein with nascent Ndc80-binding sites. Modeling was carried out for CENP-T molecules with two Ndc80 binding sites, denoted by two numbers as subscripts of the letter *T*. CENP-T has four possible Ndc80-binding configurations $T_{ij}$ (*i* refers to site 1, *j* refers to site 2): $T_{00}$, $T_{10}$, $T_{01}$, $T_{11}$. Subscript is 0 when the corresponding site is free from Ndc80, subscript is 1 when Ndc80 is bound to the corresponding site. Ndc80 binding and dissociation from the nascent (not mature) sites was described with the rate constants $k_{on}$ and $k_{off}$ (Eq. (S1)).

Maturation transition. Maturation of the Ndc80/CENP-T complex was modeled as a probabilistic event, which transforms the nascent site, where the Ndc80 molecule is already bound, into a mature site with a reduced rate of Ndc80 dissociation. Thus, maturation in the model is a first-order irreversible transition with a corresponding kinetic rate constant called the "maturation" constant ($k_{mat}$).

CENP-T protein with mature Ndc80-binding sites. Mature sites were modeled analogously to the nascent ones. Configurations of CENP-T molecules with mature sites are indicated with an asterisk: $T_{i^*j}$, $T_{ij^*}$, $T_{i^*j^*}$ $(i,j = \overline{0,1})$. The mature Ndc80/CENP-T complex dissociates very slowly, so our experiments did not provide any information about the rate of Ndc80 re-association with the mature sites. For simplicity, we assume that when both sites of CENP-T with bound Ndc80s have matured, the complex does not dissociate. Because the dissociation rate from each of the mature sites is very low, this assumption has no impact on model results. Additionally, the model includes the reaction of Ndc80 binding to mature CENP-T sites with rate constant $k_{on}$. This reaction is incorporated to provide a symmetric description of Ndc80 binding and unbinding reactions, despite the negligible concentration of mature CENP-T sites without bound Ndc80.

Within the above-described framework, the model features 13 possible configurations of CENP-T. The total number of CENP-T molecules across all configurations remains constant during the calculations and is equal to the initial number of molecules, denoted by $[T_0]$. The ratio of the number of CENP-T molecules in configuration $T_{ij}$ to $[T_0]$ is represented by $\{T_{ij}\}$, where the curly braces indicate the fraction of CENP-T in the corresponding configuration.

***Reaction scheme and equations for interactions between Ndc80 and CENP-T***

All reactions are summarized in *SI Appendix*, Fig. S7A. Transitions between different nascent sites are depicted with black arrows that are labeled with corresponding kinetic rate constants. Transitions from the intermediate (nascent) to mature forms are depicted with red (for site 1) and blue (for site 2) arrows that are labeled with corresponding maturation rate constants.

Ndc80 recruitment was modeled as a series of reversible chemical reactions between $N$ and 13 different configurations of CENP-T. For simplicity, during one step, each CENP-T molecule can bind only one molecule $N$.

<u>Binding</u> of $N$ is described as a second-order chemical reaction characterized by association rate constants $k_{1on}$ and $k_{2on}$ depending on the binding site. Same rate constants describe association to the nascent and mature sites.

<u>Dissociation</u> of $N$ from any CENP-T site is described as a first-order reaction characterized by a dissociation constant: $k_{1off}$ or $k_{2off}$ correspond to Ndc80 dissociation from the nascent sites, whereas $k^*_{1off}$ or $k^*_{2off}$ correspond to the mature sites.

<u>Maturation</u> of each site is described with maturation constant $k_{1mat}$ for site 1 and $k_{2mat}$ for site 2.

The following differential equations correspond to this reaction scheme, where *t* is time.

$$\frac{d\{T_{00}\}}{dt} = -k_{2on}[N]\{T_{00}\} + k_{2off}\{T_{01}\} - k_{1on}[N]\{T_{00}\} + k_{1off}\{T_{10}\} \quad \text{(S8)}$$

$$\frac{d\{T_{01}\}}{dt} = k_{2on}[N]\{T_{00}\} - k_{2off}\{T_{01}\} - k_{1on}[N]\{T_{01}\} + k_{1off}\{T_{11}\} - k_{2mat}\{T_{01}\} \quad \text{(S9)}$$

$$\frac{d\{T_{10}\}}{dt} = k_{1on}[N]\{T_{00}\} - k_{1off}\{T_{10}\} - k_{2on}[N]\{T_{10}\} + k_{2off}\{T_{11}\} - k_{1mat}\{T_{10}\} \quad \text{(S10)}$$

$$\frac{d\{T_{11}\}}{dt} = k_{1on}[N]\{T_{01}\} - k_{1off}\{T_{11}\} + k_{2on}[N]\{T_{10}\} - k_{2off}\{T_{11}\} - k_{1mat}\{T_{11}\} - \\ - k_{2mat}\{T_{11}\} \quad \text{(S11)}$$

$$\frac{d\{T_{0^*0}\}}{dt} = -k_{2on}[N]\{T_{0^*0}\} + k_{2off}\{T_{0^*1}\} - k_{1on}[N]\{T_{0^*0}\} + k^*_{1off}\{T_{1^*0}\} \quad \text{(S12)}$$

$$\frac{d\{T_{0^*1}\}}{dt} = k_{2on}[N]\{T_{0^*0}\} - k_{2off}\{T_{0^*1}\} - k_{1on}[N]\{T_{0^*1}\} + k^*_{1off}\{T_{1^*1}\} \quad \text{(S13)}$$

$$\frac{d\{T_{1^*0}\}}{dt} = k_{1on}[N]\{T_{0^*0}\} - k^*_{1off}\{T_{1^*0}\} - k_{2on}[N]\{T_{1^*0}\} + k_{2off}\{T_{1^*1}\} + k_{1mat}\{T_{10}\} \quad \text{(S14)}$$

$$\frac{d\,\{T_{1^*1}\}}{dt} = k_{1on}[N]\,\{T_{0^*1}\} - k^*_{1off}\{T_{1^*1}\} + k_{2on}[N]\{T_{1^*0}\} - k_{2off}\,\{T_{1^*1}\} + k_{1mat}\,\{T_{11}\} - $$
$$- k_{2mat}\,\{T_{1^*1}\} \qquad \text{(S15)}$$

$$\frac{d\,\{T_{00^*}\}}{dt} = -k_{2on}[N]\{T_{00^*}\} + k^*_{2off}\{T_{01^*}\} - k_{1on}[N]\{T_{00^*}\} + k_{1off}\{T_{10^*}\} \qquad \text{(S16)}$$

$$\frac{d\,\{T_{01^*}\}}{dt} = k_{2on}[N]\,\{T_{00^*}\} - k^*_{2off}\{T_{01^*}\} - k_{1on}[N]\,\{T_{01^*}\} + k_{1off}\,\{T_{11^*}\} + k_{2mat}\{T_{01}\} \qquad \text{(S17)}$$

$$\frac{d\{T_{10^*}\}}{dt} = k_{1on}[N]\{T_{00^*}\} - k_{1off}\{T_{10^*}\} - k_{2on}[N]\{T_{10^*}\} + k^*_{2off}\{T_{11^*}\} \qquad \text{(S18)}$$

$$\frac{d\,\{T_{11^*}\}}{dt} = k_{1on}[N]\,\{T_{01^*}\} - k_{1off}\,\{T_{11^*}\} + k_{2on}[N]\,\{T_{10^*}\} - k^*_{2off}\,\{T_{11^*}\} - k_{1mat}\{T_{11^*}\} +$$
$$+ k_{2mat}\,\{T_{11}\} \qquad \text{(S19)}$$

$$\frac{d\,\{T_{1^*1^*}\}}{dt} = k_{1mat}\,\{T_{11^*}\} + k_{2mat}\{T_{1^*1}\} \qquad \text{(S20)}$$

$$\frac{d[N]}{dt} = 0 \qquad \text{(S21)}$$

$$\frac{d}{dt}\,(\{T_{00}\} + \{T_{01}\} + \{T_{10}\} + \{T_{11}\} + \{T_{0^*0}\} + \{T_{0^*1}\} + \{T_{1^*0}\} + \{T_{1^*1}\} + \{T_{00^*}\} + \{T_{01^*}\} +$$
$$\{T_{10^*}\} + + \{T_{11^*}\} + \{T_{1^*1^*}\}) = 0 \qquad \text{(S22)}$$

The system of first-order differential equations (S8)-(S22) was used to calculate $\{T\}$ as a function of time from the start of reactions for indicated initial conditions (see below). Then, for every time point the fraction of CENP-T in each configuration was multiplied by the number of bound Ndc80 molecules for this configuration. For example: the number of bound Ndc80s to $T_{00}$, $T_{01}$ and $T_{11}$ were 0, 1 and 2 correspondingly. Changes in the sum of all fractions adjusted in this manner correspond to the kinetics of the number of Ndc80 molecules bound to one CENP-T molecule. The Ndc80 binding and unbinding (washout) stages of experiments with the coverslip-immobilized CENP-T were calculated separately.

#### *Description of the calculation algorithm*

The concentration of Ndc80 and the fractions of CENP-T molecules in all possible configurations were calculated at each iteration of the numerical solution of the system of differential equations (S8)-(S22). This system was solved using a programming and numeric computing platform MATLAB version R2020b with a versatile MATLAB ODE solver *ode45*. Ode45 is a medium order method for non-stiff differential equations that implements a Runge-Kutta method with a variable time step for efficient computation. Total calculation time was chosen to match experimental conditions.

#### *Initial conditions*

The initial concentration of Ndc80 $[N_0]$ for the binding stage was chosen to match specific experimental conditions (200 nM unless otherwise specified). Concentration of soluble

Ndc80 $[N]$ was kept constant, and the initial fraction of CENP-T molecules in all configurations was 0, except for $\{T_{00}\}$, which was set to 1 for normalization. To model the washout, soluble Ndc80 concentration was set to zero, and the initial fractions of CENP-T molecules in different configurations were taken from the results of corresponding calculations during the binding stage.

#### *Model calibration*

The model was calibrated by obtaining a good match between model predictions and experiments with different concentrations of Ndc80 Bonsai and clusters of CENP-$T^{site1}$ and CENP-$T^{site2}$ (*SI Appendix,* Fig. S8A). Since the maximum number of Ndc80 molecules that can bind to each modeled site is one, this modeling framework is not applicable to conditions where binding levels exceeded this limit. Such curves were excluded when determining binding and unbinding kinetics, but they were used for fitting the fractions of stable complexes. In the model, the association, dissociation, and maturation rate constants for a deleted site were set to zero, corresponding to the lack of any interactions with this site. Thus, constants for each site were calibrated based on the experimental data for proteins containing only this site. First, we analyzed the biphasic dissociation kinetics observed in these experiments after a 2-minute incubation with various Ndc80 concentrations and the subsequent washout of soluble Ndc80. The fast initial rate of complex dissociation determined the rate constants $k_{1off} =$ 4,000·$10^{-5}$ $s^{-1}$, $k_{2off} =$ 700·$10^{-5}$ $s^{-1}$. The second phase, corresponding to the Ndc80 dissociation from the mature complexes, was very slow, giving rise to a “stable” population. In view of the limited observation time for the stable fraction (up to 20 min), it was not possible to determine this dissociation rate accurately. Our estimates suggested similar dissociation rate for each mature site on CENP-T: $k^*_{1off} = k^*_{2off} =$ 5·$10^{-5}$ $s^{-1}$. With both dissociation rate constants fixed, the Ndc80 binding phase was then used to determine the association rate constants for each site, yielding $k_{1on}$ = 1.5·$10^{-3}$ $nM^{-1}$ $s^{-1}$ and $k_{2on}$ =4.5·$10^{-3}$ $nM^{-1}$ $s^{-1}$. The values of maturation constant for each site were chosen to ensure the best visual fit to the fractions of the stable populations: $k_{1mat} =$ 6·$10^{-4}$ $s^{-1}$ and $k_{2mat} =$ 40·$10^{-4}$ $s^{-1}$.

#### Model for soluble Ndc80 and CENP-T proteins

To predict the kinetics of complex formation in solution, the same system of equations (S8)-(S22) was used with the exception of equation (S21), which was replaced with equation (S23), as described below. Because the mass action law is directly applicable to molecular interactions in solution, the variables $\{T\}$ in this case represent soluble concentrations of the corresponding CENP-T configurations. The FCS experiments were carried out with the GFP-tagged CENP-T at 1 nM, and the unlabeled Ndc80 at 10 nM because the FCS method is only applicable to solutions with low concentration of fluorescent molecules. Thus, in the model, we could no longer assume that the concentration of soluble Ndc80 did not change during the binding reactions with CENP-T proteins. To modify the model, the concentrations of all soluble proteins were allowed to change over time according to their binding and unbinding interactions. To take this into account, equation (S21) was replaced with the following equation:

$$\frac{d[N]}{dt} = -k_{1on}[N]\{T_{00}\} - k_{2on}[N]\{T_{00}\} - k_{2on}[N]\{T_{10}\} - k_{1on}[N]\{T_{01}\} + k_{1off}\{T_{10}\} + \\ + k_{1off}\{T_{11}\} + k_{2off}\{T_{01}\} + k_{2off}\{T_{11}\} - k_{1on}[N]\{T_{0^*0}\} - k_{2on}[N]\{T_{0^*0}\} - \\ k_{2on}[N]\{T_{1^*0}\} - -k_{1on}[N]\{T_{0^*1}\} + k^*_{1off}\{T_{1^*0}\} + k^*_{1off}\{T_{1^*1}\} + k_{2off}\{T_{0^*1}\} + \\ k_{2off}\{T_{1^*1}\} - k_{1on}[N]\{T_{00^*}\} - -k_{2on}[N]\{T_{00^*}\} - k_{2on}[N]\{T_{10^*}\} - k_{1on}[N]\{T_{01^*}\} + \\ k_{1off}\{T_{10^*}\} + k_{1off}\{T_{11^*}\} + +k^*_{2off}\{T_{01^*}\} + k^*_{2off}\{T_{11^*}\} \qquad \text{(S23)}$$

In line with the conditions of the FCS experiments, the model was used to analyze the binding phase only (no washout). The initial concentration of CENP-T $[T_0]$ was 1 nM, concentration of Ndc80 $[N_0]$ was 10 nM. Model results for changes in concentrations of selected CENP-T configurations were normalized to the initial concentration of CENP-T molecules to present the predictions as fractions of the formed complexes, enabling direct comparison with the experimental outcomes.

For the CENP-T$^{2D}$ protein, the plotted fraction corresponds to configurations with two bound Ndc80 molecules ($T_{11}$, $T_{1*1}$, $T_{11*}$, $T_{1*1*}$). For mutant CENP-T proteins with only one site, the plotted fractions correspond to complexes with one Ndc80 molecule. To build concentration dependencies, the initial concentration of Ndc80 $[N_0]$ was varied in the range from 0 to 400 nM and the fraction on Ndc80/CENP-T complexes was determined after 70 min interaction.

To generate predictions in the absence of site maturation, same model was used but the maturation rate constants for both sites were set to 0. Three different CENP-T proteins were modeled: CENP-T$^{2D}$, CENP-T$^{site1}$ and CENP-T$^{site2}$. The values of association rate constants remained the same as in the model with maturation for the corresponding proteins, as listed in *SI Appendix,* Table S1. The kinetics of Ndc80 binding to CENP-T in solution was then calculated using two sets of values of the dissociation rate constants for the corresponding proteins: for the nascent and mature sites.

**Summary of results**

<u>Interaction between soluble Ndc80 protein and immobilized clusters with CENP-T$^{site1}$ and CENP-T$^{site2}$</u>

The experiments involving a 2-minute incubation of these clusters with Ndc80, followed by washout, were used to calibrate the model, resulting in the rate constants listed in the *SI Appendix,* Table S1 (see section 'Model calibration'). With these values of rate constants, the model describes reasonably well the kinetic curves for a range of soluble Ndc80 concentration (*SI Appendix,* Fig. S8A,B). These rate constants were then applied to describe our other experimental results with CENP-T site1 and site2 protein clusters. Specifically, the model was used to calculate the number of stably bound Ndc80 molecules after variable incubation time with 200 nM Ndc80, followed by the washout. A good match was obtained with no adjustment to any model parameters (*SI Appendix,* Fig. S8C). Thus, quantitative analysis of the results with clusters containing proteins with only one CENP-T site revealed that the kinetic rates for their intermediate complexes with Ndc80 (i.e. for nascent sites) are highly similar, with site 2 having 3-fold faster association rate and ~ 6-fold slower dissociation. Additionally, site 2 matures ~7-fold faster, which makes site 2 overall "stronger" than site 1 (faster recruitment of strongly bound Ndc80 protein). However, once Ndc80 forms strong binding interfaces with any of these sites, the mature complexes of site 1 and site 2 are equally stable up to the maximum duration of our experiments.

<u>Interaction between soluble Ndc80 protein and immobilized clusters with CENP-T$^{2D}$</u>

The same set of constants was applied to experiments with immobilized CENP-T$^{2D}$ clusters, which contain both binding sites. If the properties of these sites do not depend on the presence of the second site, the constants determined for individual sites should also provide a good fit to the results of experiments with CENP-T$^{2D}$ clusters, which contain CENP-T with both Ndc80 binding sites and the same phosphomimetic substitutions as in the mutant site1 and site2 proteins. Indeed, the kinetic constants for formation of intermediate complexes with both sites within the CENP-T$^{2D}$ clusters, as well as stability of the mature complexes, were adequate. However, both sites matured faster in the context of the CENP-T$^{2D}$ clusters than within the clusters with single site proteins. To obtain a better match with the results with CENP-T$^{2D}$ clusters, the maturation rate constant for site 1 was increased 3.3-fold, while the maturation rate for site 2 was increased 7.5-fold (*SI Appendix,* Fig. S8C, Table S1). Thus, maturation rates of both sites are accelerated in the presence of each other. However, the rate of maturation for site 2 remains significantly faster than for site 1.

Some of our experiments used clusters of CENP-T$^{6D}$ protein, which in addition to the two phosphomimetic substitutions with aspartic acids in each of the two binding sites (T11D and T85D) present in the CENP-T$^{2D}$ protein, has four additional substitutions: T27D, S47D, T195D, S201D. The experimental results with this protein were highly similar to those with CENP$^{2D}$, although formation of stable complexes within the CENP-T$^{6D}$ was slightly slower (*SI Appendix,* Fig. S6E). Accordingly, formation of the stable Ndc80/CENP-T complexes in CENP-T$^{6D}$ clusters incubated with 200 nM Ndc80 were well described using the same constants as for CENP-T$^{2D}$, but a slightly better match was obtained using the 4-fold reduction of the maturation rate for site 1 in the CENP-T$^{6D}$ clusters than in the CENP-T$^{2D}$ clusters (*SI Appendix,* Fig. S8C). With this modification, the concentration dependency for CENP-T$^{6D}$ clusters and 1-200 nM Ndc80 for 2 min interaction time was also matched (*SI Appendix,* Fig. S8D). Thus, it appears that some (or all) of these additional phosphorylations (T27D, S47D, T195D, S201D) reduce sensitivity of CENP-T to its molecular environment.

<u>Interaction between soluble Ndc80 protein and immobilized CENP-T monomers</u>

Our model makes no assumptions about the nature of Ndc80 interactions with monomers versus clusters of CENP-T, assuming identical molecular mechanisms. If Ndc80 interacts with monomeric CENP-T at the same rate constants as with CENP-T in clusters, the model predictions should be the same for both molecular forms of CENP-T, as the same system of equations describes the underlying chemical reactions. However, applying the model to experimental results for monomers of the same CENP-T species yielded a poor fit because the fraction of stable Ndc80/CENP-T complexes is much lower in monomers versus clusters. In the model, the rate of accumulation of stable complexes strongly depends on the rate of maturation transition. Thus, a much better fit was achieved using adjusted maturation rates, as described for each type of CENP-T protein. The results for both monomeric proteins CENP-T$^{2D}$ and CENP-T$^{6D}$ were well fit using the same kinetic constants for the formation of intermediate complexes and the dissociation rate of the mature complexes as used for the clusters containing these proteins. However, the maturation rate for both proteins was decreased 3-fold and 38-fold for site 1 and site 2, respectively (*SI Appendix,* Fig. S9A,B, Table S1). The results for monomeric proteins containing only one of the sites (CENP-T$^{site1}$ and CENP-T$^{site2}$) were well fit using the same dissociation constants that were found to fit the results of experiments with clusters containing these respective proteins. The association constant for site2 also provided an adequate fit, but for CENP-T protein containing site 1, a slightly better fit was achieved by reducing its association rate constant 3-fold. Importantly, to achieve a good fit, the maturation rate for both proteins had to be decreased: 6-fold for site1 and 8-fold for site2 (*SI Appendix,* Fig. S9C,D, Table S1).

<u>Interaction between soluble Ndc80 and CENP-T proteins</u>

The model with parameter values for CENP-T monomers (*SI Appendix,* Table S1) was applied to describe results of the FCS experiments in which 1 nM CENP-T (CENP-T$^{2D}$, CENP-T$^{site1}$ or CENP-T$^{site2}$) was incubated with 10 nM Ndc80. The fraction of complexes was calculated for 0-90 min time range and plotted using a 1 s time step together with experimental data points (Fig. 1J). For all CENP-T proteins, the maturation-dependent binding kinetics in solution is predicted to have the initial phase with a fast increase in complex formation, corresponding to transient interactions with the nascent sites and formation of the intermediate complexes. This initial fast phase is followed by a slower phase, which arises owing to improved Ndc80 retention. As alternative models, we calculated predictions for the maturation-free binding with two dissociation rates $k_{off}$ and $k_{off}^{*}$, corresponding to intermediate and stable complexes. Unlike the model with maturation, these control models predict a simple exponential increase in the number of Ndc80/CENP-T complexes followed by a $k_{off}$-dependent plateau (broken and dotted lines in Fig. 1J). Because the experimental curves feature bi-phasic kinetics, the experimental data align closely with the predictions derived from the model with maturing sites, rather than models with no maturation. The best fit for the maturation model was with the same constants as in *SI Appendix,* Table S1, except for the association constants, which had to be slightly modified: $k_{1on}$ was increased 3-fold in CENP-T$^{2D}$ and 4-fold in CENP-T$^{site1}$ soluble monomers, whereas $k_{2on}$ was decreased 1.8-fold for both CENP-T$^{2D}$ and CENP-T$^{site2}$ soluble monomers. These adjustments relative to the same monomers attached to the glass surface may reflect a different accessibility of CENP-T sites to Ndc80 molecules in solution vs. the coverslip surface. Additionally, the model with maturation matches results of experiments in which 1 nM CENP-T was incubated for 70 min with the range of Ndc80 concentrations (0-400 nM with 1 nM step) without any change in model parameters (Fig. 1K). This remarkable consistency between model and experiment strongly indicates that the two-step binding mechanism is a bona fide feature of Ndc80-CENP-T interactions and is not an artifact of the TIRF-based imaging assay.

**Model limitations and conclusions**

Our chemical kinetics model is based on fundamental principles of chemical kinetics, and therefore, it cannot provide insights into the molecular mechanisms of complex formation or dissociation. However, given the extensive experimental data we have collected using various CENP-T proteins and different methodologies, the model is essential for testing the consistency across these diverse results. We focused on reactions occurring on the coverslip surface, ensuring the applicability of chemical kinetics modeling to this context. Additionally, we designed the model's output in terms of molecule count rather than concentration, allowing for direct comparison with experimental data.

Our approach involved calibrating the model using a specific dataset, enabling the identification of parameter values that provided a reasonable fit. Same parameters and equations were then applied to other experimental datasets, allowing us to identify which constants needed adjustment for a closer alignment with experimental results. Notably, the goal of this approach was not to achieve a quantitatively perfect match between the model and experiments; thus, the reported constants should be regarded as estimates. This approximation more accurately reflects the experimental uncertainty in determining exact values, which often vary across experimental repetitions. While the derived constants may shift if a different set of experiments is used for calibration, the major differences and their ratios remain robust.

Importantly, the conclusion that the Ndc80 reaction follows a two-step mechanism does not rely on the model or its calibration methods. Instead, it is derived entirely from model-independent data analysis. With this in mind, we summarize the key outcomes of our modeling approach.

First, the model allows us to estimate the affinity of Ndc80 binding to CENP-T. According to *SI Appendix,* Table S1, within the CENP-T monomer, the affinity of the intermediate complex formed at site 1 is 25 nM, which is approximately 10-fold lower than the affinity of the Ndc80-CENP-T complex at site 2 (1–2 nM). The rate of maturation into a stable complex at site 1 is 4-fold slower than at site 2 ($2 \times 10^{-4}$ vs. $8 \times 10^{-4}$ $s^{-1}$). Thus, site 1 is overall weaker, leading to a slower accumulation of stable complexes with Ndc80 compared to site 2. However, once the stable complex is formed, both sites appear equally stable, though further methods may resolve subtle differences in the future.

Second, a wide range of our experiments using different CENP-T monomers could be explained by these kinetic constants. This consistency across diverse approaches enhances confidence in the conclusions drawn from our data. We note, however, that minor adjustment of certain constants sometimes improved the model's fit. For instance, changes in association rate constants—such as a 3-fold increase for site 1 in solution versus on the coverslip surface, and a three-fold decrease for this site within the CENP-$T^{site1}$ protein—may reflect the sensitivity of site 1 to its specific environment. Nevertheless, these small differences may not be significant due to experimental and modeling limitations. Similarly, maturation of both sites appears faster when both are present in CENP-$T^{2D}$, compared to proteins containing only one site. These differences do not exceed two-fold and should not be overstated.

Third, experiments using different CENP-T clusters could be well described using the same binding/unbinding rate constants as for their respective monomers. However, for all proteins and experiments, the maturation rates that worked well for the monomers had to be increased when CENP-T proteins were clustered. For site 2, the maturation rate in CENP-$T^{2D}$ clusters increased nearly 40-fold, while clustering CENP-$T^{site2}$ led to a 8-fold increase. This suggests that both sites are highly sensitive to their molecular environment, with site 2 being particularly responsive and additionally influenced by the presence of site 1. Interestingly, many observed effects between individual sites and proteins containing both sites are not additive and do not follow simple rules, a pattern also seen in mutant proteins with "composite" sites. Therefore, the exact molecular mechanism governing the transition between intermediate and stable Ndc80/CENP-T complexes, as well as the enhancement effects of a dense molecular environment, remains to be fully understood.

**Note 3. Structural analysis of human CENP-T sites bound to Spc24/25 head**

The crystal structure of the chicken CENP-T region (63-93 aa; T72D, S88D) in complex with the Spc24/25 head has provided valuable insights into the molecular details of this interaction (19). The central α-helix of the CENP-T region fits snugly into a groove on the Spc24/25 surface that faces the outer kinetochore (*SI Appendix* Fig. S10D, E). The inserted helix appears to function as a key structural element supporting the load-bearing tethering of the Ndc80 complex. It is "suspended" by the C-terminal flanking region, which exits perpendicularly from the groove and wraps around the Spc24/25 head, ultimately connecting to the centromere via the extended CENP-T shaft. The N-terminal flank makes a sharp turn away from the helix, with this configuration being regulated by phosphorylation at the critical threonine residue T72 (19). This N-terminal region lies on a rugged surface of Spc24, forming a third binding interface within this intricate topology. The overall configuration resembles the S-wrap used in rope climbing.

The structure of human Ndc80/CENP-T complex has not yet been solved. The N-terminus of human CENP-T has two regions with high sequence homology to the chicken Ndc80-binding site: site 1 (1-30 aa) and site 2 (76-106 aa). Both sites contain conserved sequence for Cdk- phosphorylation at T11 and T85 (Fig. 3A), which enhances human CENP-T binding to Ndc80 (19, 26). For our structural analyses, we employed CENP-T sequences with the phosphomimetic substitutions at these sites.

To predict the binding configuration between human CENP-T sites and Ndc80, we employed the AlphaFold software (16, 18), see Materials and Methods section ‘Modeling of Ndc80/CENP-T complexes using AlphaFold software’. To validate this approach, we predicted structures for Spc24(134-195 aa)/Spc25(134-232 aa) head of the chicken Ndc80 complex, which had been crystallized previously (PDB: 3VZ9) (19). Structural similarity between the predicted and known structures was quantified using the Template Modeling (TM) score, which equals 1 for the perfect match (21, 22). The TM score for chicken Spc24/25 structures was 0.98 (*SI Appendix*, Fig. S11A). The TM score for predicted chicken Spc24/Spc25 head complexed with a fragment of chicken CENP-T (63-93 aa with T72D and S88D substitutions) and the analogous structure PDB:3VZA (19) was 0.98 (*SI Appendix*, Fig. S11B). Lastly, the AlphaFold2 prediction for the human Spc24(122-197 aa)/Spc25(118-224 aa) head yielded a TM score of 0.91 when aligned with the corresponding X-ray structure in the Ndc80 Bonsai complex (PDB: 2VE7) (20) (*SI Appendix*, Fig. S11C). These high scores suggest that AlphaFold software is a reliable tool for investigating the structural conformations of the Spc24/25 head in complex with human CENP-T sites.

The enhanced retention of Ndc80 after the saturation of both CENP-T binding sites strongly implies that upon Ndc80 association, the sites change their molecular properties. However, the molecular origins of the different binding kinetics by these sites are unclear. We noticed a marked similarity in the secondary structure of human CENP-T sites relative to the chicken CENP-T complexed with their cognate Spc24/25 proteins (*SI Appendix*, Fig. S10E). The position of this central α-helix of CENP-T in all three sites appears to be highly conserved. The local Distance Difference Test (pLDDT) indicated high level of confidence for the predictions of central α-helical regions (*SI Appendix*, Fig. S12A). For site 1, the high confidence (pLDDT > 90) extended from 11 to 21 aa, whereas for site 2 it spanned from 86 to 104 aa. To probe reproducibility of these predictions, we performed five independent AlphaFold simulations for each pair of proteins and quantified their differences using standard deviations (SDs) of each Cα atom position (*SI Appendix,* Fig. S12B). These metrics indicate that the central α-helix maintains a conserved position across human sites, residing within the groove of the Spc24/25 subunits (*SI Appendix*, Movies S1 and S2).

These analyses also show that the C-terminal flanking region (C-flank) following the α-helix at human site 2 exhibits a configuration similar to that of the C-flank in chicken CENP-T, wrapping around the Spc25 subunit (*SI Appendix*, Fig. S10E). The human C-flank at site 2 displays a high confidence score (pLDDT > 90), low variability, and an increased number of contacts with the Spc24/25 head (*SI Appendix,* Fig. S12), indicating stable association. In contrast, the N-terminal flanking region (N-flank) of human site 2 has a much lower confidence score. However, both human site 2 and chicken CENP-T share a conserved leucine residue (L81 in humans, L68 in chickens) (*SI Appendix*, Fig. S10C). This leucine has been shown to play a crucial role in hydrophobic interactions with the Spc24 subunit (19). Thus, configuration of human site 2 with the Spc24/Spc25 head has similar tripartite organization.

Furthermore, there were notable differences between the flanking regions of site 1 and site 2 (Fig. 3B; *SI Appendix*, Movies S1 and S2). The average confidence score for the C-flanking region of site 1 was only ~ 55 (*SI Appendix*, Fig. S12D), significantly lower than the confidence score for the same region in site 2 (~90). Additionally, the predictions for the C-flank of site 1 exhibited higher standard deviations (SD) (*SI Appendix*, Fig. S12B) and lower number of contacts with Spc24/25, relative to site 2 (*SI Appendix*, Fig. S12C). The N-flanking region of site 1 was also predicted with low confidence (~40) and it lacks the conserved hydrophobic leucine residue (*SI Appendix*, Fig. S10C).

Together, these results suggest that the distinct kinetics of human site 1 and site 2 binding to Spc24/25, as well as their differing rates of maturation, may arise from variations in the structure and composition of their flanking regions. These differences likely influence how the regions wrap around the globular heads. To further explore the molecular determinants of these differing maturation rates, we conducted AlphaFold simulations on CENP-T peptides with composite sites: flanks2/helix1 and flanks1/helix2 (*SI Appendix*, Movies S1 and S2). Notably, the flanking regions largely maintained their characteristic configurations, even when paired with the heterologous helix (Fig. 3D). This effect was particularly pronounced in the behavior of the C-terminal flanking region, which displayed low confidence and high variability in the flanks1/helix2 peptide, mirroring its behavior with helix1. Conversely, the C-flanks of the flanks2/helix1 peptide remained highly organized, consistent with its behavior alongside helix2. Therefore, sequences of the flanking regions likely play an important role in defining their dynamic behavior. Because AlphaFold does not directly assess protein dynamics, these findings do not exclude the possibility that the helical regions themselves are crucial in driving the maturation transition. Another limitation of AlphaFold-based modeling is its focus on sequences containing binding sites, without accounting for other unstructured regions of CENP-T, which may also contribute to the process. Consequently, future studies employing methods that probe conformational dynamics are necessary to fully elucidate the molecular basis of the maturation transition.

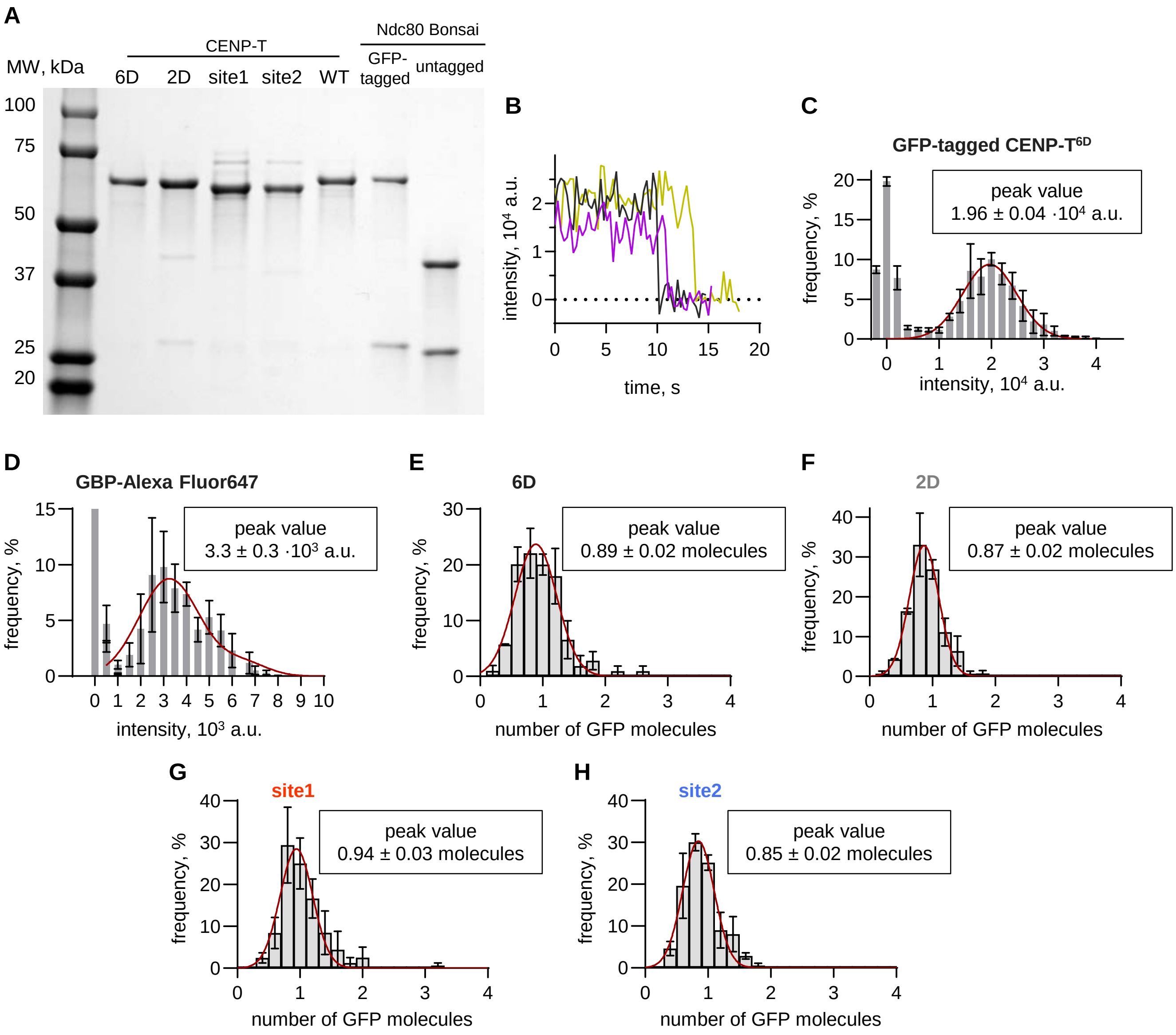


**Fig. S1. Characterization of GFP-CENP-T monomers.**

**(A)** Purified CENP-T and Ndc80 constructs used in this study were analyzed by SDS-PAGE. **(B)** Example photobleaching curves for coverslip-immobilized GFP-tagged CENP-$T^{6D}$ dots. **(C)** Histogram of integral intensities collected from photobleaching curves for GFP fluorophore, number of independent experiments (N) = 3, total number of analyzed dots (n) = 47. Binned intensities are represented with mean ± SEM, red lines are fittings of the main distributions with Gaussian functions. Peaks of intensities close to zero correspond to background values. **(D)** Same as in panel (C) but for GBP-Alexa Fluor 647, N = 4, n = 112. **(E-H)** Histograms of the number of GFP molecules per fluorescent dot in the chamber with coverslip-immobilized indicated GFP-tagged CENP-T proteins, N = 3, n > 100 per protein. Peak values correspond to average number of CENP-T molecules per fluorescent dot in our experiments. For all proteins, dots containing several GFP molecules were rare and they likely resulted from two or more molecules localizing close together.

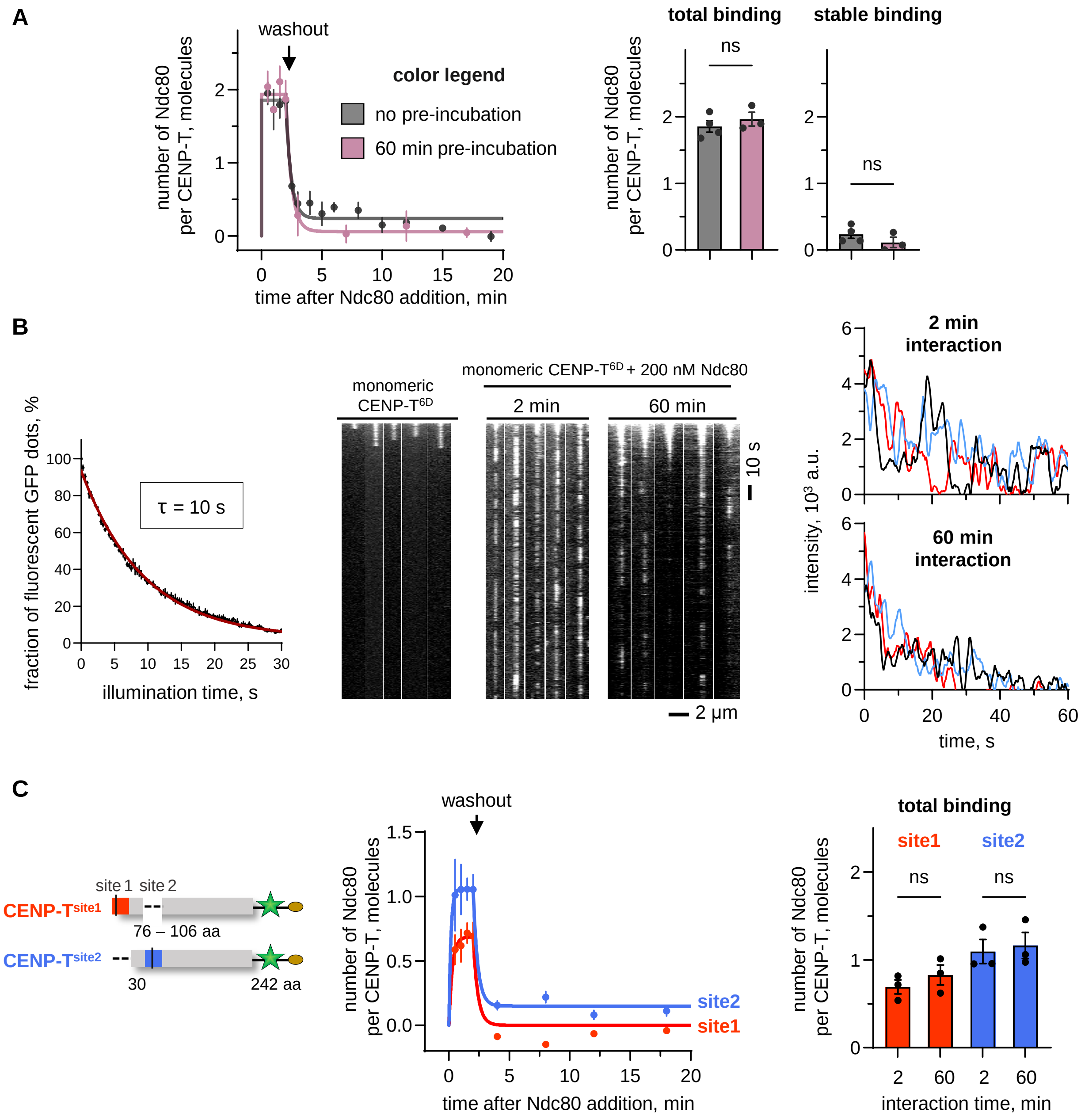


**Fig. S2. Interaction between Ndc80 and different CENP-T monomers.**

**(A)** Left: Ndc80 (200 nM) interaction with coverslip-immobilized CENP-T$^{6D}$ monomer. Soluble Ndc80 was added immediately after immobilization of CENP-T$^{6D}$ molecules (no pre-incubation, grey) or after 60 min pre-incubation of immobilized CENP-T$^{6D}$ molecules on the coverslip (60 min pre-incubation, pink). Lines are exponential fittings; each point is mean ± SEM, N = 3-4. Panels on the right show results for total and stable Ndc80 binding in these experiments. Each point represents median determined in independent kinetic experiment with n > 20 molecules per each time point; bars show mean ± SEM; unpaired t-test with Welch's correction: ns = p > 0.05. Data for CENP-T$^{6D}$ without pre-incubation are the same as in Fig. 1G,H. **(B)** Graph on the left shows fraction of fluorescent GFP-CENP-$T^{6D}$ dots as a function of illumination time. Each point represents the mean ± SEM from N = 3. Kymographs show photobleaching of immobilized monomeric CENP-$T^{6D}$ in the absence and presence of Ndc80-GFP at different time since the start of incubation. Graphs on the right show representative line scans for the kymographs, illustrating reduced signal after longer interaction time between CENP-T and Ndc80. **(C)** As in panel (A) but showing Ndc80 interaction with CENP-T monomers containing only one of the binding sites.

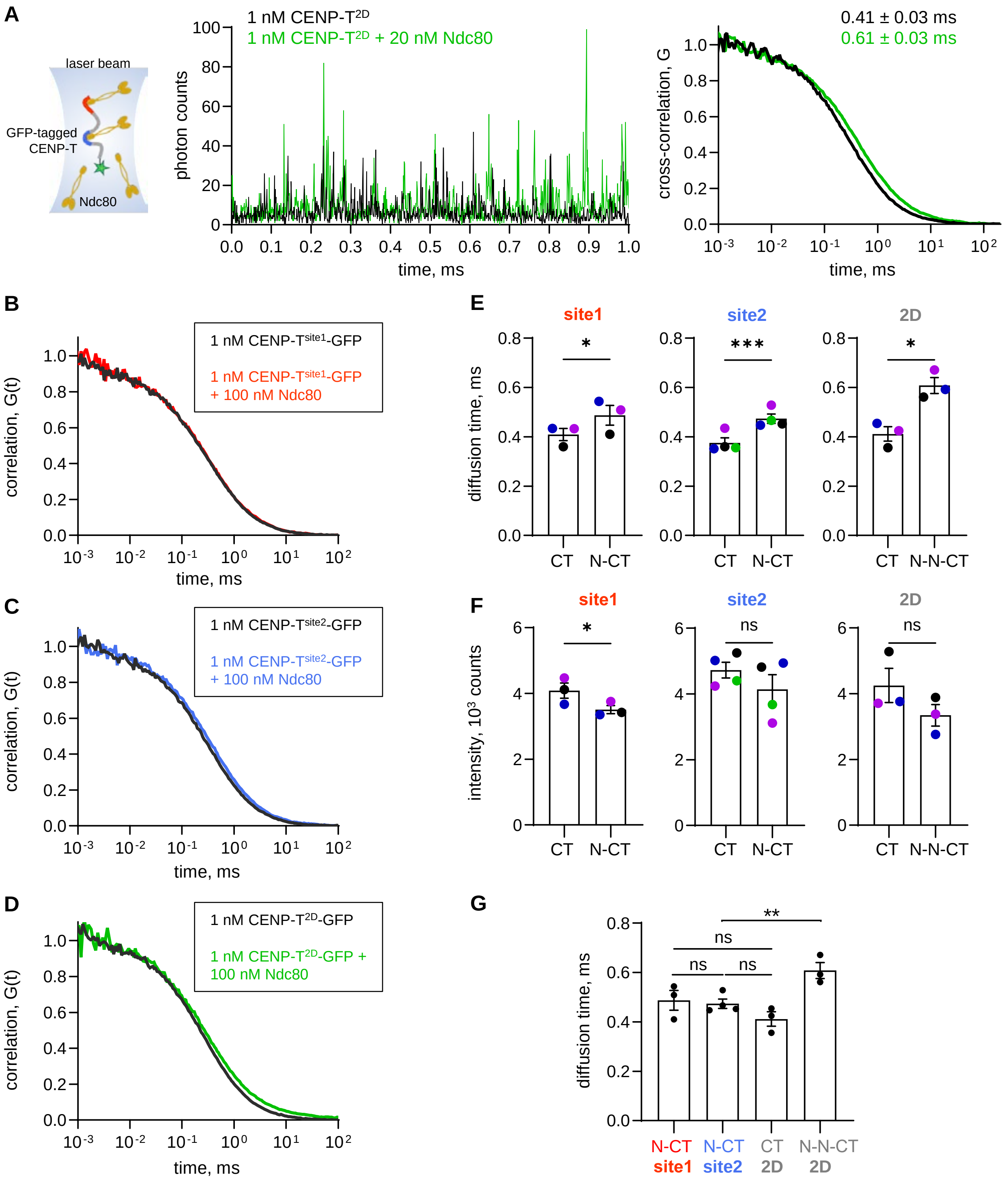


**Fig. S3. Analysis of interaction between soluble Ndc80 and CENP-T using FCS approach.**

**(A)** Scheme of FCS assay, example time traces, and corresponding cross-correlation curves for 1 nM GFP-tagged CENP-T$^{2D}$ alone or with 20 nM unlabeled Ndc80. Shift between the cross-correlation curves indicates change in the diffusion times (numbers about the curves) due to complex formation. **(B)-(D)** Example cross-correlation curves for indicated GFP-tagged CENP-T constructs alone or in the presence of 100 nM unlabeled Ndc80. **(E)** The diffusion time and **(F)** fluorescence intensity of the indicated CENP-T (CT) proteins alone or in a complex with one or two Ndc80 molecules (N-CT and N-N-CT). Each point represents an independent experiment, and points of the same color represent experiments carried out concurrently. Statistical significance was determined with a paired t-test: ns = $p > 0.05$, * = $p < 0.05$, *** = $p < 0.001$. **(G)** Diffusion times of indicated CENP-T complexes alone or with Ndc80 molecules. Statistical significance was determined using unpaired t-test with Welch's correction: ns = $p > 0.05$, ** = $p < 0.01$.

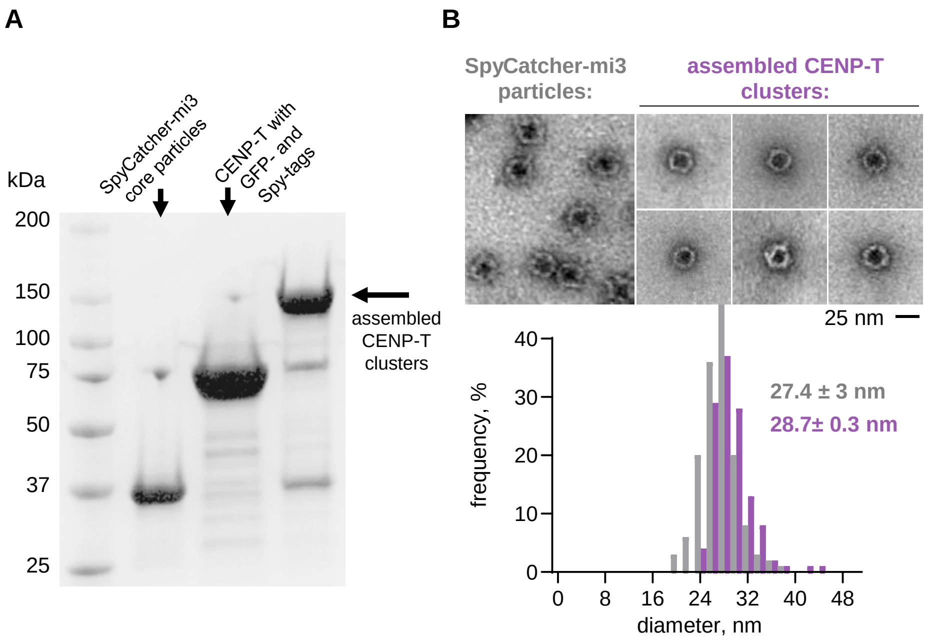


**Fig. S4. Preparation and electron microscopy of CENP-T clusters.**

**(A)** SDS-PAGE for conjugation of CENP-T$^{2D,short}$ with SpyCatcher-mi3 particles. **(B)** Example images of SpyCatcher-mi3 core particles before and after conjugation to CENP-T visualized by negative staining electron microscopy. Histogram of the diameter of SpyCatcher-mi3 core particles before (grey columns) and after conjugation to CENP-T (purple columns), based on n = 140 SpyCatcher-mi3 core particles from N = 4 experiments, and n = 120 assembled CENP-T clusters from N = 3 experiments. Although the conjugated CENP-T is not resolved, there is a slight increase in particle size.

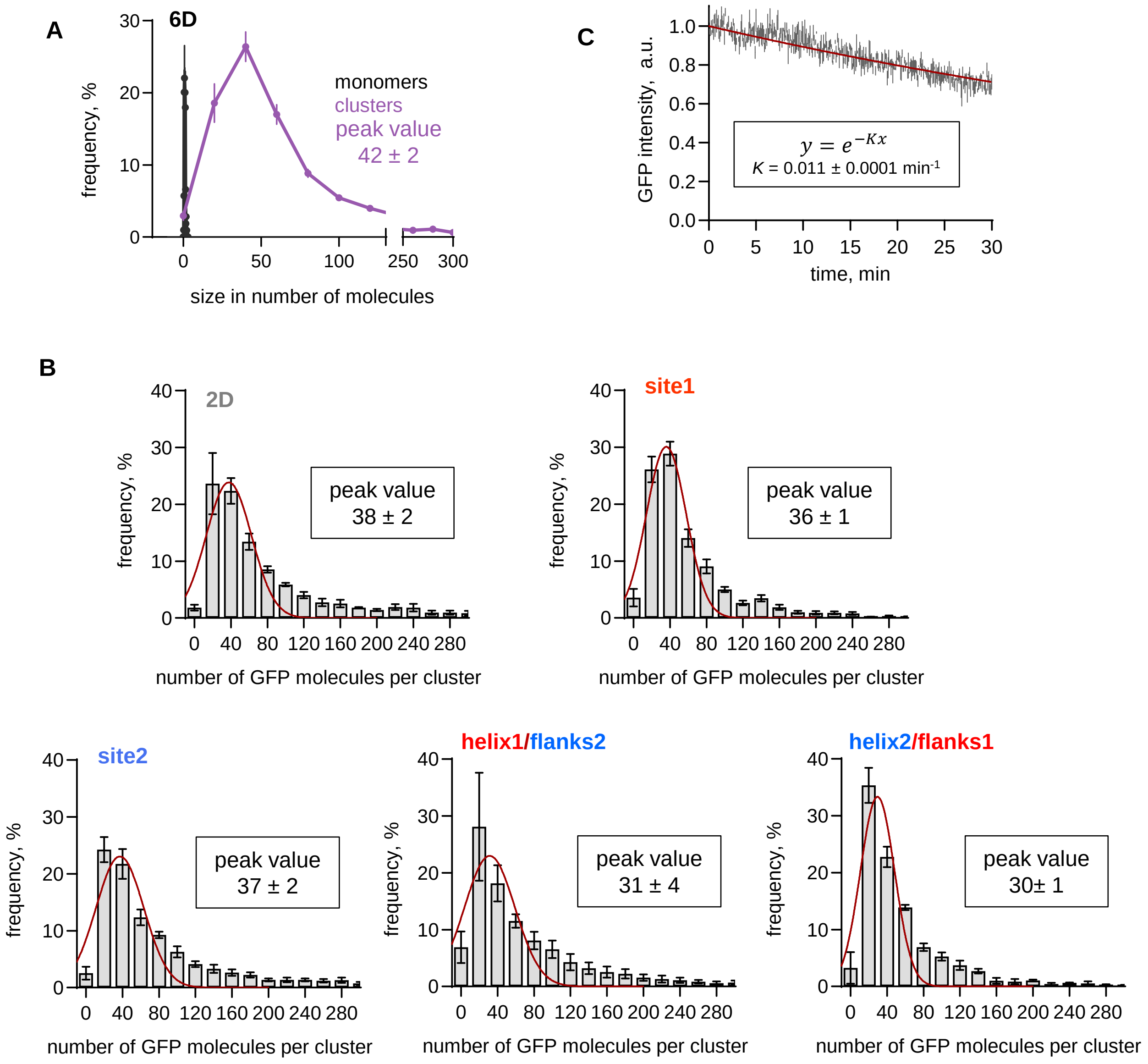


**Fig. S5. Determining size of the CENP-T clusters.**

**(A)** Histogram of the number of GFP molecules per cluster of the GFP-tagged CENP-T$^{6D}$ protein plotted together with the histogram for GFP-tagged CENP-T$^{6D}$ monomers (same as in ***SI Appendix***, **Fig. S1E**). Each bin shows mean ± SEM from N = 3-24 experiments with n > 150 clusters each. **(B)** Column histograms showing cluster size as in panel (A). Numbers are peak values determined with Gaussian fitting (red line), corresponding to average number of CENP-T molecules per cluster. **(C)** Intensity during photobleaching of clusters with GFP-tagged CENP-T with exponential fitting.

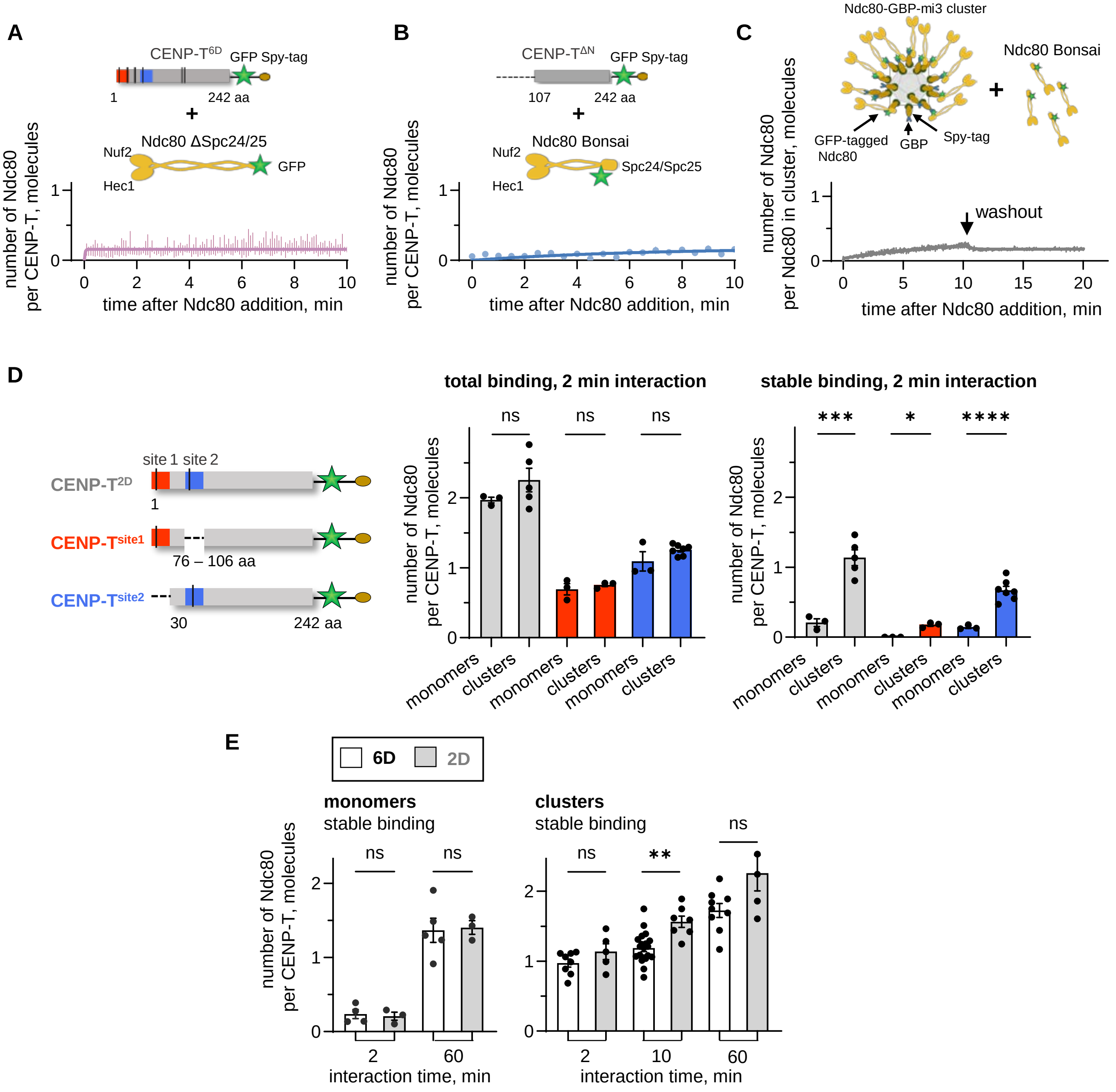


**Fig. S6. Interaction between Ndc80 and different protein clusters.**

(A) Binding of Ndc80 ΔSpc24/25 (200 nM) to clustered CENP-T$^{6D}$. Soluble Ndc80 was added at time 0; line id exponential fit. (B) Same as in panel (A) but using Ndc80 Bonsai (200 nM) and clustered CENP-T$^{\Delta N}$, which lacks the N-terminus with Ndc80 binding sites. (C) Same as in panel (A) but using Ndc80 Bonsai and Ndc80-GBP-mi3 clusters for binding and washout. (D) Ndc80 Bonsai (200 nM) interaction (2 min) with clusters containing CENP-T$^{2D}$, CENP-T$^{site1}$ or CENP-T$^{site2}$ and their respective monomers. Each point represents an independent experiment with n > 14 clusters/molecules; bars show the mean ± SEM; unpaired t-test with Welch's correction: ns = $p > 0.05$, * = $p < 0.05$, *** = $p < 0.001$, **** = $p < 0.0001$. (E) Stable Ndc80 binding to monomeric and clustered CENP-T$^{6D}$ vs. CENP-T$^{2D}$. Each point represents an independent experiment with 200 nM Ndc80 for n > 12 monomers/clusters; bars show the mean ± SEM; unpaired t-test with Welch's correction: ns = $p > 0.05$, ** = $p < 0.01$

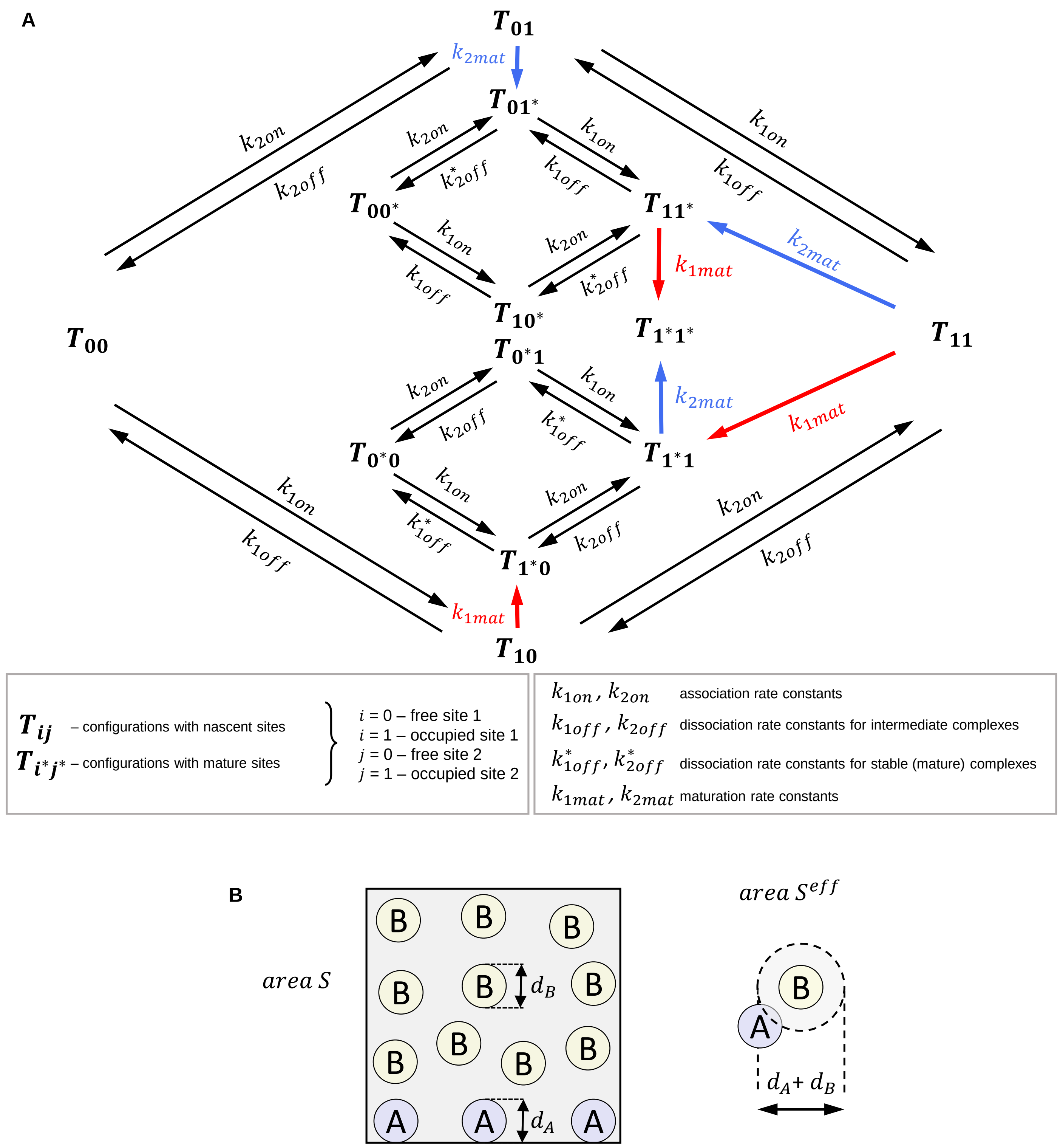


**Fig. S7. Mathematical model of Ndc80 interaction with CENP-T binding sites.**

**(A)** The reaction scheme depicting kinetic reactions between CENP-T (T) and Ndc80 (N) molecules. Arrows forming the outer contour represent the initial binding and unbinding of Ndc80 molecules to nascent sites on CENP-T, so they correspond to intermediate complex formation/dissociation. The red and blue arrows correspond to maturation transitions and lead to the reaction arrows forming the inner contours, corresponding to Ndc80 interaction with mature sites (indicated with asterisk). **(B)** Schematic illustrating interaction areas for immobilized molecules *A* (diameter $d_A$) and soluble molecules *B* (diameter $d_B$).

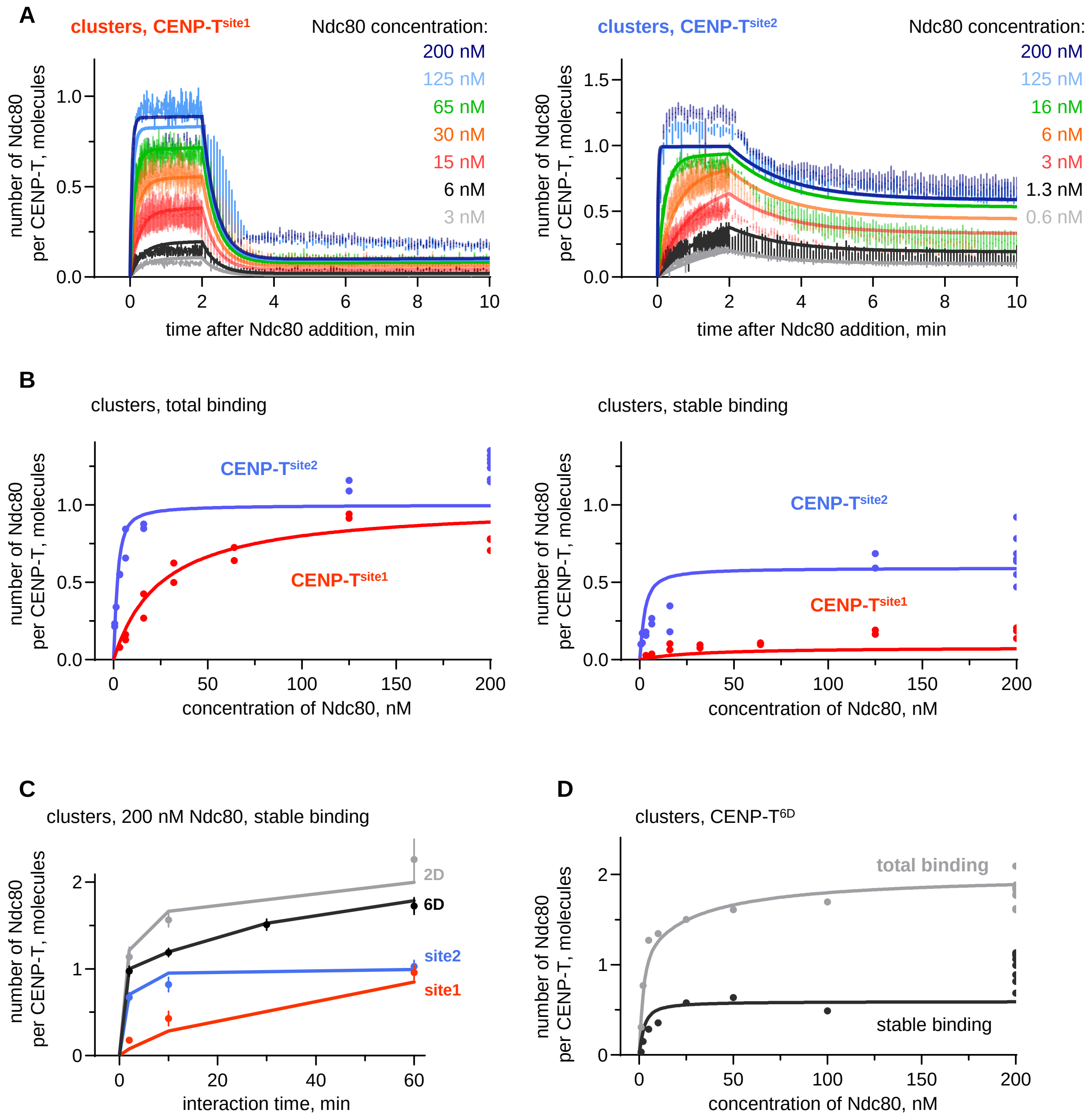


**Fig. S8. Model results for interaction between soluble Ndc80 and CENP-T clusters.**

Graphs show theoretical predictions (lines) overlaid with experimental results (vertical ticks and points). **(A)** Binding and unbinding of Ndc80 Bonsai to clusters with CENP-T$^{site1}$ or CENP-T$^{site2}$ proteins. Vertical ticks show mean with SEM from N = 2-7 experiments for different Ndc80 concentration (color coded). **(B)** Total and stable binding of Ndc80 Bonsai to indicated CENP-T clusters as a function of Ndc80 concentration after 2-min interaction. Each point represents an independent experiment with n > 12 clusters. **(C)** Stable binding of Ndc80 Bonsai (200 nM) to indicated CENP-T clusters as function of interaction time. Each point is mean ± SEM from N = 2-20 experiments. **(D)** As in panel (B) but for clusters containing CENP-T$^{6D}$.

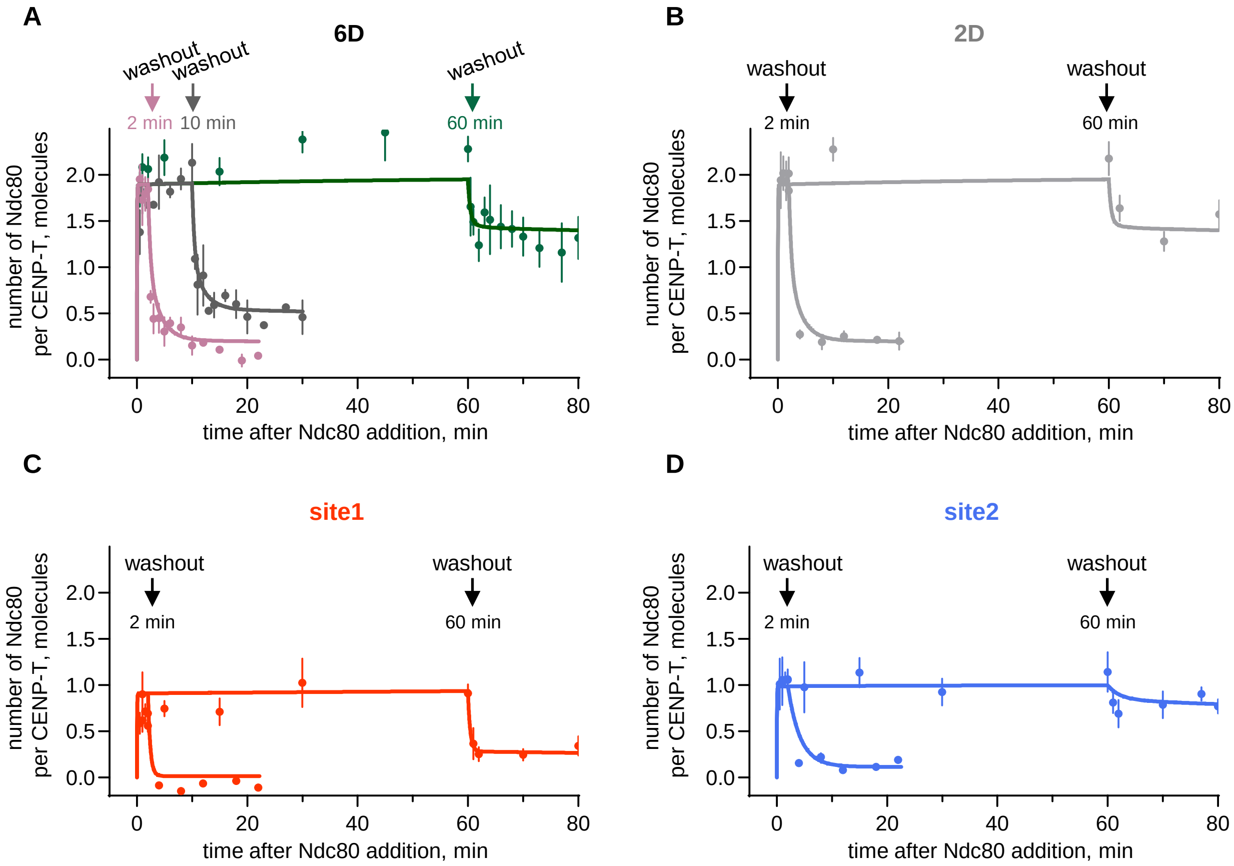


**Fig. S9. Model results for interaction between soluble Ndc80 interaction and CENP-T monomers.**

Graphs show theoretical predictions (lines) and experimental results (data points) for interaction between immobilized CENP-T monomers and soluble Ndc80 Bonsai (200 nM) added at time 0 and removed by washout at indicated times. Experimental results for CENP-T$^{6D}$, CENP-T$^{site1}$ and CENP-T$^{site2}$ are the same as in Fig. 1F,G and *SI Appendix* Fig. S2C. For CENP-T$^{2D}$ data points are mean ± SEM from N = 3 experiments.

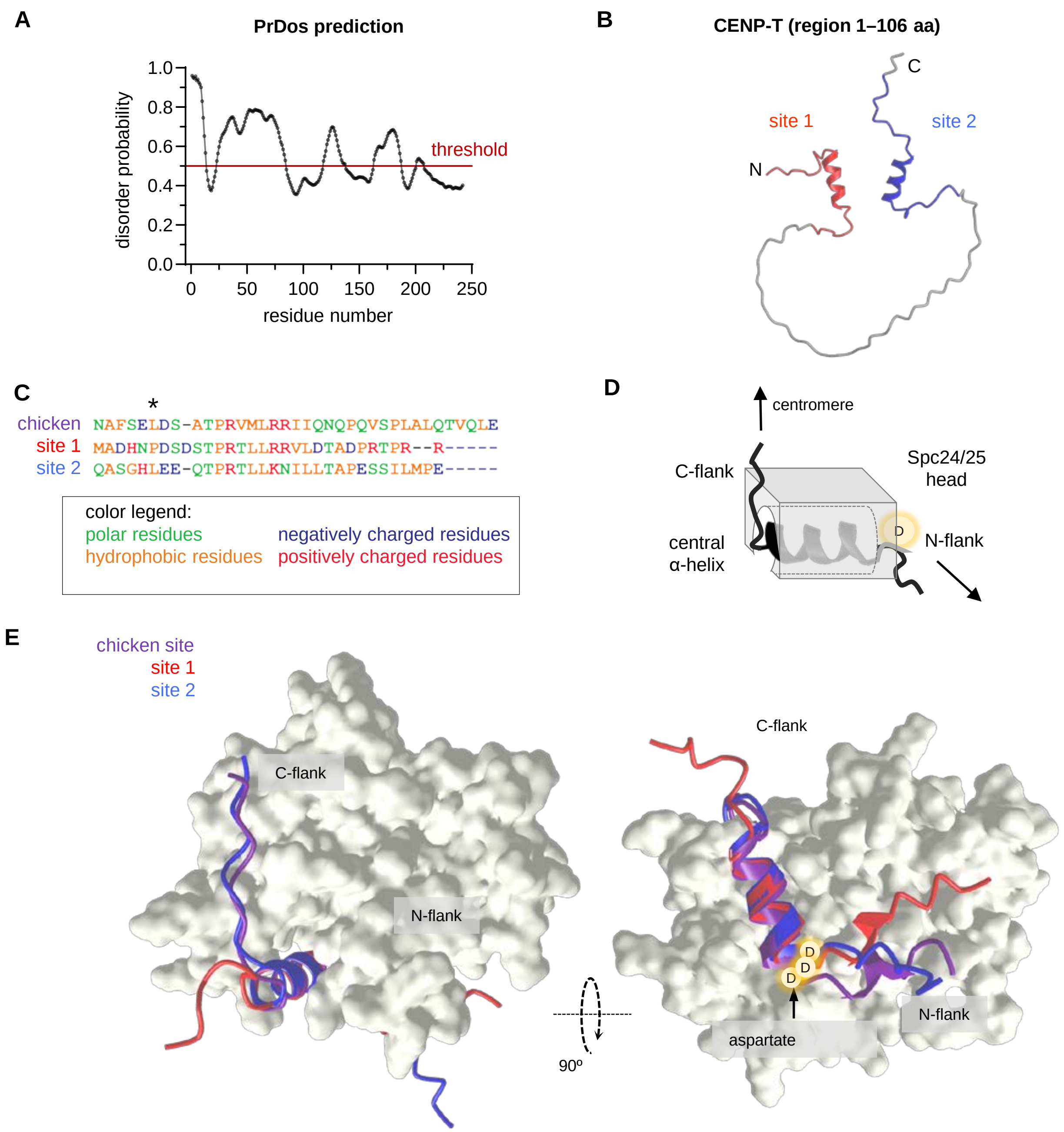


**Fig. S10. Structural features of chicken and human CENP-T binding sites.**

**(A)** Disorder probability of human CENP-T N-terminal region (1-242 aa) predicted by PrDOS software. Amino acids with a disorder probability above 0.5 are expected to be disordered with false positive rate 5%. **(B)** Structure of N-terminal region of CENP-T (1-110 aa) as predicted by AlphaFold2. Regions corresponding to site 1 (1-30 aa) are shown in red, and for site 2 (76-106 aa) in blue. **(C)** Aligned sequences of chicken CENP-T's site (63-93 aa), human CENP-T site 1 (1-30 aa) and site 2 (76-106 aa) showing differences in charge and hydrophobicity. Within the sequences, essential leucine and its proline replacement in site 1 is indicated with * symbol. **(D)** A simplified schematic of the 30 aa peptide of CENP-T (binding site) forming an S-wrap around the Spc24/25 head based on 3vzA structure for chicken proteins. **(E)** Crystal structure of chicken CENP-T's site in complex with Spc24/25 (PDB: 3VZA, (19)) aligned with the configurations of human sites 1 and site 2 as predicted by AlphaFold2.

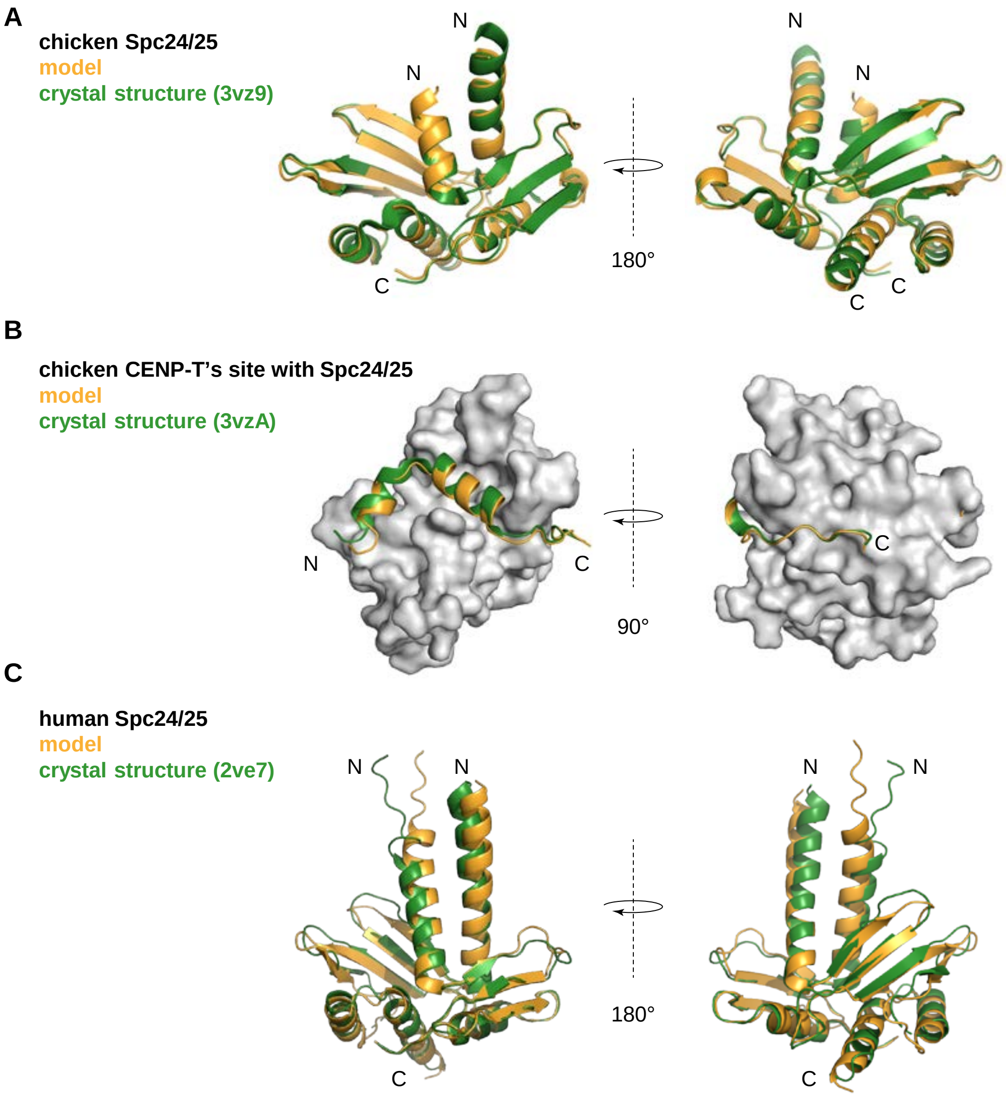


**Fig. S11. Verification of AlphaFold predictions for Spc24/25 domains.**

**(A)** The crystal structure of chicken Spc24(134-195 aa)/Spc25(134-232 aa) complex (PDB: 3vz9;(19, 20)) is aligned with the structure of the same proteins predicted by AlphaFold2. **(B)** Crystal structure of chicken Spc24(134-195aa)/Spc25(134-232 aa) in a complex with phosphomimetic CENP-T (63-93 aa; T72D, S88D) (PDB: 3vza; (19)) is aligned with the structure of the same proteins predicted by AlphaFold2. **(C)** Crystal structure of human Spc24(122-197 aa)/Spc25(118-224 aa) (PDB: 2ve7; (20)) is aligned with the structure of the same proteins predicted by AlphaFold2.

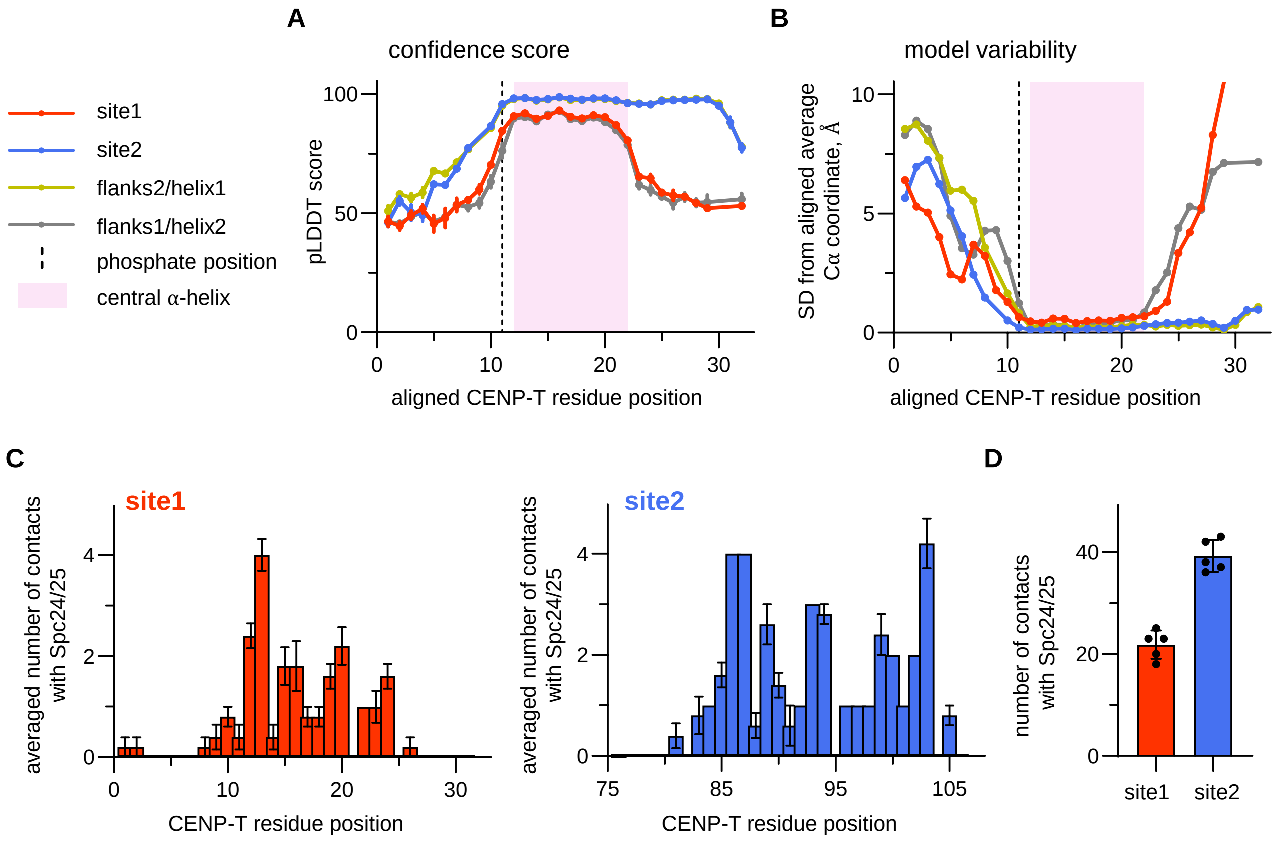


**Fig. S12. AlphaFold-based analysis of binding human CENP-T to Spc24/25 head**.

(**A**) The plot shows pLDDT score (mean ± SEM) for the Cα atoms of the CENP-T chain in the complex between Spc24/25 and indicated CENP-T sites for N = 5 predictions of AlphaFold3. Score 100 corresponds to maximal confidence. Different sequences were aligned to match position of the critical conserved phosphate. Green trace is not visible when its overlays with the blue trace. (**B**) The curves show standard deviations (SD) of the Cα coordinates of each amino acid in N = 5 predictions of AlphaFold3. Larger SD corresponds to more variable configurations. **(C)** Histograms show the average number of amino acid contacts between the Spc24/25 head and CENP-T peptides corresponding to site 1 or 2. Each bin shows mean ± SEM from N = 5 predictions of AlphaFold3. **(D)** Graph shows the total number of contacts (mean ± SD) made by site 1 and site 2 with Spc24/25 head for N = 5 AlphaFold3 simulations.

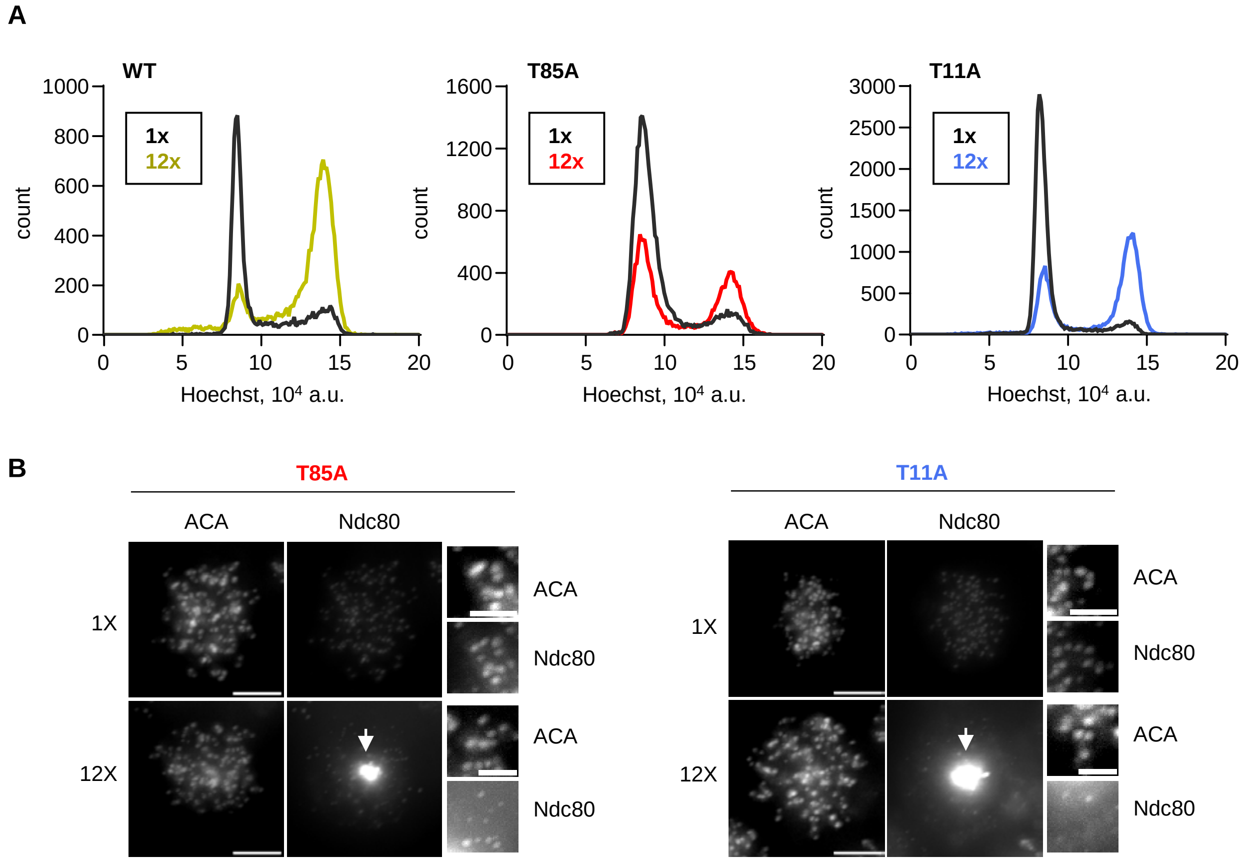


**Fig. S13. SunTag-based approach to study effect CENP-T oligomerization in cells.**

**(A)** Histograms showing the distribution of DNA content stained with Hoechst in HeLa cells expressing the indicated CENP-T constructs, which are either monomeric (1x) or oligomerized using the SunTag system into 12x-oligomers. **(B)** Representative images of mitotic cells showing localization of kinetochores and CENP-T mutant oligomers with the indicated number of repeats (white arrows). The centromeres (ACA) and Ndc80 were visualized with immunofluorescence. Images with different CENP-T constructs have different image adjustments, and the insets are adjusted differently from the full-size images for improved visibility. Scale bars: 5 µm.

**Table S1. Kinetic rate constants for interactions between Ndc80 and CENP-T proteins in different molecular forms**

The table provides values of the rate constants that provided visually good fit for experiments using TIRF microscopy, as described in section “Summary of results”, see also section “Reaction scheme and equations for interactions between Ndc80 and CENP-T”. Briefly, $k_{on}$ - association rate constant; $k_{off}$ – dissociation rate constants for different complexes (intermediate complex with a nascent site and stable complex with a mature site); $k_{mat}$ – maturation rate constant; subscript 1 stands for site 1, subscript 2 for site 2, superscript * indicates constants for mature sites.

| rate constant | site | units | CENP-T$^{2D}$ | | CENP-T$^{site1}$ | | CENP-T$^{site2}$ | |
|---|---|---|---|---|---|---|---|---|
| | | | monomers | clusters | monomers | clusters | monomers | clusters |
| $k_{1on}$ | site 1 | $10^{-3}$ $nM^{-1}$ $s^{-1}$ | 1.5 | 1.5 | 0.5 | 1.5 | 0 | 0 |
| $k_{2on}$ | site 2 | $10^{-3}$ $nM^{-1}$ $s^{-1}$ | 4.5 | 4.5 | 0 | 0 | 4.5 | 4.5 |
| $k_{1off}$ | nascent site 1 | $10^{-5}$ $s^{-1}$ | 4,000 | 4,000 | 4,000 | 4,000 | 0 | 0 |
| $k_{2off}$ | nascent site 2 | $10^{-5}$ $s^{-1}$ | 700 | 700 | 0 | 0 | 700 | 700 |
| $k^*_{1off}$ | mature site 1 | $10^{-5}$ $s^{-1}$ | 5 | 5 | 5 | 5 | 0 | 0 |
| $k^*_{2off}$ | mature site 2 | $10^{-5}$ $s^{-1}$ | 5 | 5 | 0 | 0 | 5 | 5 |
| $k_{1mat}$ | | $10^{-4}$ $s^{-1}$ | 2 | 20 | 1 | 6 | 0 | 0 |
| $k_{2mat}$ | | $10^{-4}$ $s^{-1}$ | 8 | 300 | 0 | 0 | 5 | 40 |

### Legends to Movies

#### Structure of human Spc24/25 head with bound peptides corresponding to wild type and composite CENP-T sites.

Structures were predicted using AlphaFold3 software and the best scoring models from five independent simulations are shown. The human Spc24/25 head, composed of Spc24 (134-195 aa) and Spc25 (118-224 aa), is shown in grey. The corresponding static structures are depicted in Figure 3 panels B and D. Positions of the phosphomimetic substitution are shown in yellow in each CENP-T peptide. Degrees for the corresponding rotations are shown in the left upper corner; C – C-termini, N – N-termini.

**Movie S1.** Predictions for CENP-T site 1 (1-30 aa), note the disordered flanking regions.

**Movie S2.** Predictions for CENP-T site 2 (76-106 aa). The configuration of the C-terminal flanking region is more consistent, with this flank aligns closely along the surface of the Sp24/25 head.

**Movie S3.** Predictions for the composite CENP-T site 2, containing flanking regions of original site 2 and helix of site 1. C-terminal flanking region aligns along the surface of the Spc24/25 head, as in the original site 2. The configuration of predicted N-terminal region is variable and appears to follow a different pattern than in the original site 2.

**Movie S4.** Predictions for the composite CENP-T site 2, containing flanking regions of site 1 and original helix of site 2. The unstructured flanking regions point away from the Spc24/25 head, similarly to their configuration in the original site 1 (see Movie S1).
